# Directed evolution of a SelB variant that does not require a SECIS element for function

**DOI:** 10.1101/2025.02.04.636165

**Authors:** Satoshi Ishida, Arno Gundlach, Clayton W. Kosonocky, Andrew D. Ellington

**Affiliations:** Department of Molecular Biosciences, University of Texas at Austin, Austin, TX, USA

## Abstract

In bacteria the incorporation of selenocysteine is achieved through the interaction of the selenocysteine specific elongation factor (SelB) with selenocysteine-charged tRNA^Sec^ and a selenocysteine insertion sequence (SECIS) element adjacent to an opal stop codon in a mRNA. The more generalized, SECIS-independent incorporation of selenocysteine is of interest because of the high nucleophilicity of selenium and the greater durability of diselenide bonds. It is likely that during the course of evolution selenocysteine insertion originally arose without the presence of a SECIS element, relying only on SelB. Herein we undertake experiments to evolve an ancestral version of SelB that is SECIS-independent and show that not only can this protein (SelB-v2) generally incorporate selenocysteine across from stop codons, but also that the new, orthogonal translation factor can be repurposed to other amino acids, such as serine. Given the delicate energetic balancing act already performed by EF-Tu, this achievement raises the possibility that greatly expanded genetic codes that relied in part on SelB-based loading can now be contrived.

## Introduction

In *E. coli*, the selenocysteine (Sec) incorporation machinery is coded in the genome as SelABC^1,2^. The tRNA^Sec^ (SelC) is charged by seryl tRNA synthetase, and selenocysteine synthase (SelA) converts the charged serine into selenocysteine^3,4^. SelB is particularly important as it is an elongation factor that specifically recruits Sec-tRNA^Sec^ to the ribosome, loading it adjacent to a specific RNA structure, the SECIS [selenocysteine insertion sequence] element, that is found only near a few opal (UGA) codons^5^.

In addition, the unique properties of Sec (higher nucleophilicity compared to cysteine; higher stability of diselenide bond compared to disulfide bond) makes this ‘21’st amino acid’ a potentially useful addition to an expanded genetic code^6,7^. Rather than site-specifically incorporating Sec adjacent to the SECIS, previous efforts have attempted to broaden incorporation by engineering a Sec-tRNA^Sec^ variant that is capable of being recognized by the more general endogenous elongation factor, EF-Tu. Söll and colleagues achieved this by transplanting regions of tRNA^Sec^ into tRNA^Ser8^ and further engineering through rational design yielding tRNA^UTuX9^, while Thyer and colleagues achieved this by directed evolution of anti-determinant sequences within tRNA^Sec^, making it recognizable by EF-Tu (yielding tRNASec^UX^)^10^. We propose to achieve similar outcomes by engineering a SECIS-independent SelB.

The engineering of SelB to be SECIS-independent is also an interesting prospect since it might then serve as an ‘orthogonal’ analogue of EF-Tu. EF-Tu loads tRNAs charged with their correct canonical amino acids in part by balancing the affinity to the individual tRNA (especially the T-stem) in one set of interactions with the affinity to the amino acid in another set of interactions ^11–14^. An expanded genetic code is likely to either not fit into or upset this balance, or both, given that new amino acids would be unlikely to have balanced affinities with their tRNA partners. Researchers have taken on this problem by optimizing the EF-Tu amino acid binding pocket for new amino acids such as phosphoserine or phosphotyrosine^15,16^. However, given the inherent difficulty of EF-Tu’s balancing, having an orthogonal EF-Tu functioning in parallel would allow a more ready expansion of the code than constantly changing and rebalancing the endogenous EF-Tu.

During the initial adoption of the nominal 21^st^ amino acid, it seems likely that SelB evolved from EF-Tu, and was further constrained by the SECIS element to avoid misincorporation of selenocysteine. We therefore propose to return to this ancestral state by generating a SelB variant that is not dependent upon the SECIS. A SECIS-independent SelB would not only open a new path to the production of selenoproteins, but would be a generally manipulable orthogonal translation factor for loading other amino acids. Herein, we demonstrate that a SECIS- independent SelB can in fact robustly produce selenoproteins without a SECIS, and that it can be further manipulated for the incorporation of another amino acid, serine.

## Results

### Designing SelB to function without its SECIS

The crystal structure of full length SelB in complex with a ribosome loaded with a mRNA-bearing SECIS element and Sec-tRNA^Sec^ has been solved^17^. This structure reveals that binding of SelB to the SECIS essentially tethers Sec-tRNA^Sec^ to the ribosome, with few direct interactions with the ribosome except between the C-terminal domain of SelB, helix h16 of 16S ribosomal RNA (rRNA), and protein S4. In contrast, EF-Tu is recruited by interacting directly with the L7/12 stalk of the 50S ribosomal subunit^18^. Previously mutating interactions between SelB and the SECIS resulted in a 600-fold reduction of GTPase activity^19,20^, suggesting that the SECIS:SelB interaction was functionally important.

This structure proved invaluable for designing SelB variants that might operate independently of the SECIS^17,21^. Given that the N-terminal region (Domains I to III) of SelB is a homologue of EF-Tu, while the C-terminal domain (Domain IV) consists of four winged-helix (WH) motif, wherein the last two WHs directly interact with the SECIS element^19,21,22^, our first hypothesis was that we might obtain SECIS independence through the simple expedient of deleting WH3 and WH4. This was achieved by removing WH3/WH4 (S488 to K614). This join cleanly deletes the C-terminal 381 residues and leaves a SelB variant that is only 487 residues long (**Fig. 1**).

**Figure 1.**
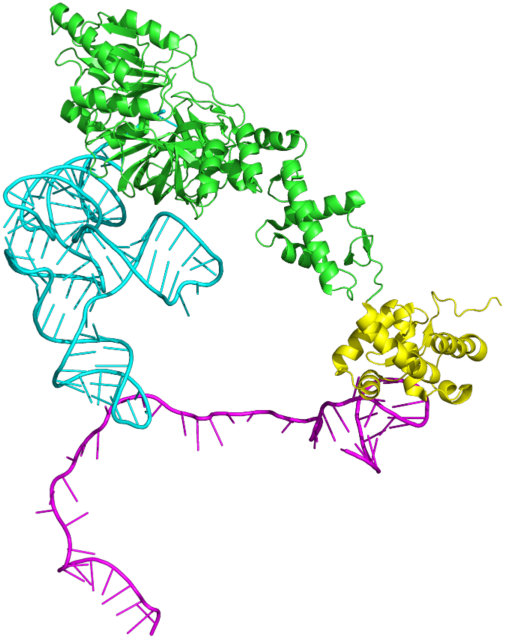
Crystal structure of SelB in complex with tRNA^Sec^ and SECIS. The purple region is the SECIS element, while the green and yellow regions are portions of the full length SelB; the yellow region is the C-terminal domain that was truncated in this study. Cyan is tRNASec. PDB ID: 5lzd.

### Assaying truncated SelB for selenocysteine incorporation

Previously, we have manipulated the machinery for selenocysteine incorporation in C321dA, an ‘Amberless’ *E. coli* strain^23^. The SelABC gene cluster was further deleted from the genome of this strain, and the necessary components for selenocysteine incorporation were instead re- introduced on a plasmid^10^ (**Fig. 2a**). In particular, SelA, SelC, were constitutively expressed, while SelB was placed under the control of an araBAD promoter.

**Figure 2.**
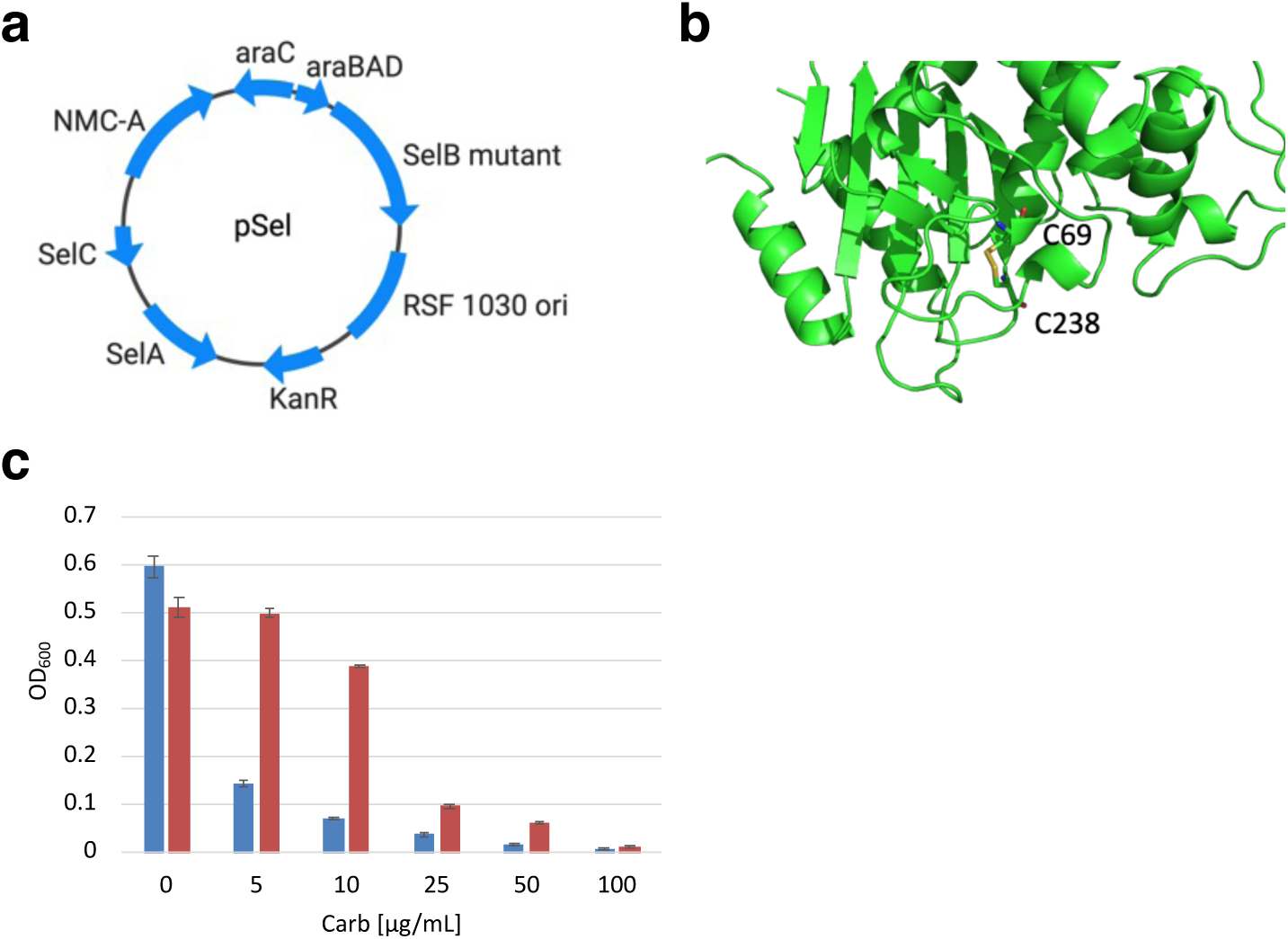
Initial activity of wild-type SelB versus truncated SelB (**a**) Map of the plasmid used for SelB expression. The actual sequence of plasmid is shown in **Supplementary Table 3**. SelA and selC are constitutively expressed while SelB is under the control of araBAD promoter. (**b**) Crystal structure of NMC-A showing the essential disulfide bond between C69 and C238 (PDB: 1BUE). (**c**) Growth in the presence of carbenicillin following induction of either WT-SelB (blue) or truncated-SelB (red). Each data point represents the mean of three independent experiments ± s.d (standard deviation).

In order to assay and select for selenocysteine incorporation, we had also previously replaced a key cysteine residue in a disulfide-dependent beta-lactamase, NMC-A, with selenocysteine (**Fig. 2b**; ^10,24^). Formation of a chimeric cysteine-selenocysteine bond promotes growth in the presence of carbenicillin. Surprisingly, the truncated SelB variant actually showed improved Sec incorporation compared to the wild-type enzyme, at least in the NMC-A assay (**Fig. 2c**). These results hint at a different role for the SECIS than is typically reported^20^: the SECIS does not enable incorporation, but rather limits mis-incorporation.

### Directed evolution of the SECIS-independent SelB variant

The demonstrated dependence of selenocysteine incorporation for NMC-A beta-lactamase activity enables the directed evolution of Sec incorporation machinery (**Fig. 3a**). Since it wasn’t clear exactly what portions of truncated SelB needed to be optimized, a library of SelB variants was generated by error-prone PCR with a mutation rate of up to 12 substitutions per gene. The transformation efficiency in the first and subsequent rounds of selection always exceeded 10^6. The first round of selection was conducted at 5 μg/mL carbenicillin, followed by three rounds of selection at 10 μg/mL carbenicillin (**Supplementary Table 1**). Error prone PCR was conducted after Round 2, with a mutation frequency of an additional 10 mutations per gene.

**Figure 3.**
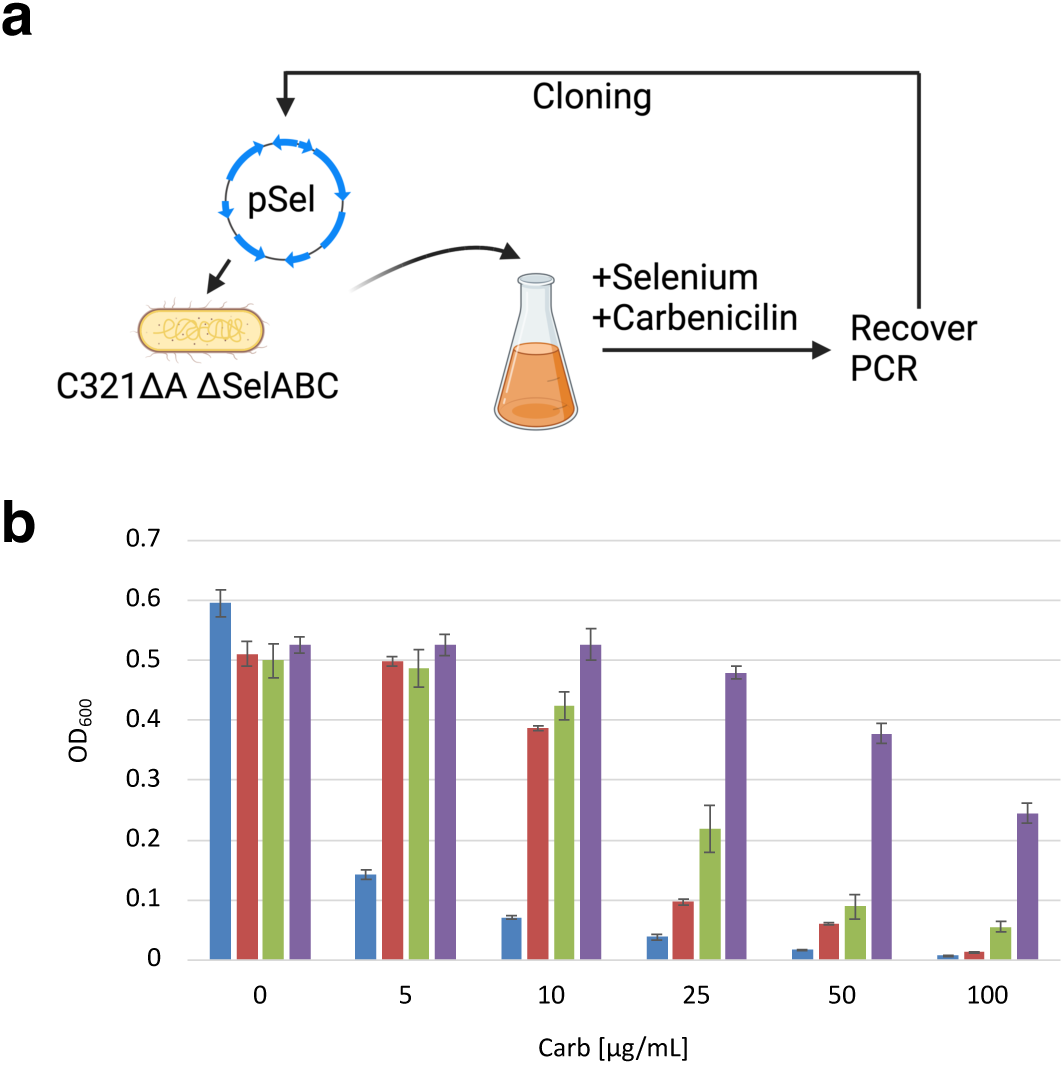
Directed evolution of SelB (**a**) Schematic of the directed evolution of SelB. A plasmid-based SelB library was transformed into C321 dSelABC and grown in the presence of both carbenicillin and selenium. Grown cells were collected and plasmid was recovered, then SelB variants were amplified by PCR and cloned back into the same backbone. A detailed summary of the rounds of selection is shown in **Supplementary Table 1**. (**b**) Growth assay in the presence of cabenicillin with WT-SelB (blue), truncated SelB (red), SelB-v1 (green) and SelB-v2 (purple). Each data point represents the mean of three independent experiments ± s.d (standard deviation).

After four rounds of selection, six individual colonies were picked (at 100 μg/mL carbenicillin), and sequencing revealed a predominant (5/6) SelB variant that shared a total of eleven substitutions, including three silent mutations (**Supplementary Table 2**). At the same time, we found that the plasmid backbone had duplicated the NMC-A gene, and that a recombination event had yielded a SelA-SelB fusion protein. These additional modifications to the plasmid should both have led to the ability to survive the carbenicillin challenge: additional copies of the NMC-A gene would yield greater beta-lactamase activity, while the fusion led to constitutive production of SelB, as opposed to the inducible araBAD promoter (**Supplementary Table 5** and **Supplementary** Fig. 2).

The enriched SelB variant was re-cloned to the original plasmid configuration, yielding SelB-v1. This plasmid was then tested for beta-lactamase activity, sans gene duplications and fusions. Sec incorporation was indeed enhanced at 100 μg/mL carbenicillin compared to truncated SelB (**Fig. 3b**).

SelB-v1 was subjected to additional rounds of directed evolution. The library for selection was prepared by error-prone PCR of SelB-v1, which contained 11 mutations including three silent mutations (**Supplementary Table 2**), and had a mutation frequency of up to 4 mutations per SelB gene. In order to increase the stringency for selenocysteine incorporation, two TAG codons were introduced into the NMC-A gene, which reduced NMC-A expression (as observed by carbenicillin resistance; **Supplementary** Fig. 3). Six additional rounds of selection were carried out at 10 μg/mL carbenicillin, and a final round at 100 μg/mL (**Supplementary Table 1**). Error prone PCR was also conducted after Rounds 3 and 5.

Some 10 colonies were picked for sequencing: all contained E463K, and 9 out of 10 contained E246K. The addition of these two predominant mutations to SelB-v1 yields SelB-v2. SelB-v2 showed enhanced incorporation efficiency compared to SelB-v1 via the NMC-A assay with increasing carbenicillin concentrations (**Fig. 3b**).

### Validating selenocysteine incorporation with SelB-v2

In order to further validate selenocysteine incorporation by SelB-v2, we attempted to introduce a fluorescent protein that depends on the incorporation of Sec for its fluorescence (smURFP^25^) into the C321dA strain. The smURFP protein covalently binds the chromophore biliverdin (BV), whose production requires the expression of the pbsA1 heme oxidase for chromophore maturation. However, transformation and expression of this pathway proved difficult, partially due to the growth defects associated with C321dA^26–28^.

Therefore, we switched to the more commonly used *E. coli* cloning and expression strain DH10B; this switch should also provide additional evidence of the generality of the utility of SelB-v2 for general selenocysteine suppression. The genomic SelABC gene in DH10B was removed to avoid competition with the native selenocysteine insertion machinery.

The fluorescence of the smURFP reporter was rendered dependent upon selenocysteine incorporation machinery by converting the cysteine codon at amino acid position 62 to TAG) ^25^. For this selenocysteine-dependent smURFP construct in the modified DH10B strain, fluorescence was only observed when (i) SelB-v2 (not wild-type SelB) was induced, and (ii) selenium was added to the media (**Fig. 4a**). In addition, for a sfGFP reporter bearing a TAG codon at position 150, fluorescence could only be observed when SelB-v2 was induced and selenium was added to the media (**Fig. 4b**).

**Figure 4.**
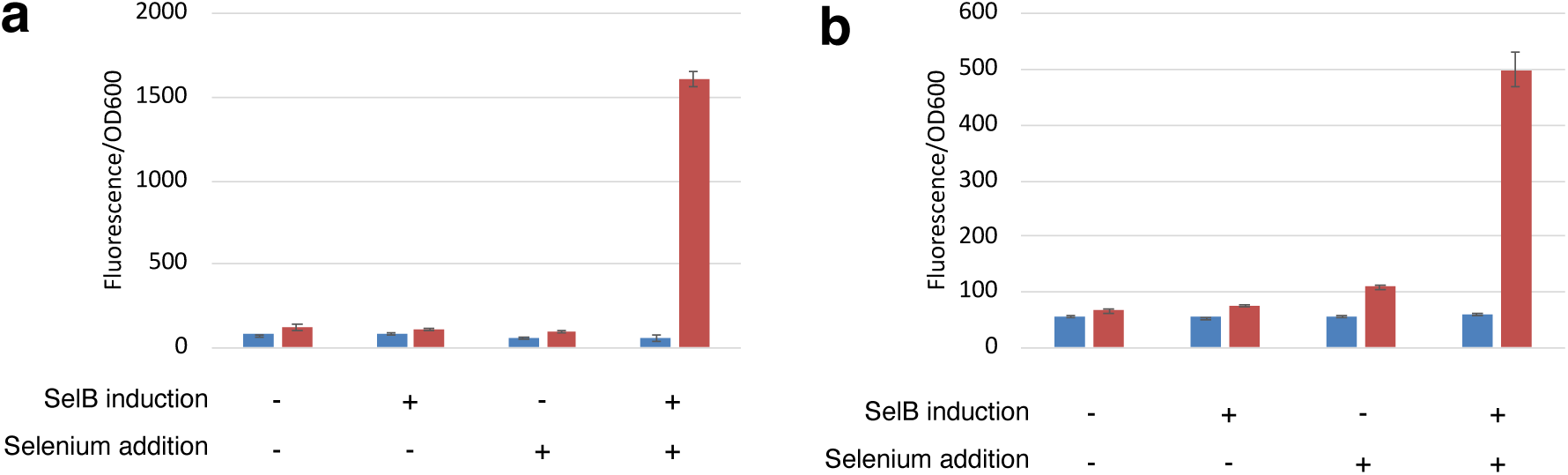
Monitoring Sec incorporation with fluorescent proteins (**a**) smURFP expression in the presence or absence of expression of SelB WT or SelB-v2, and in the presence or absence of the addition of selenium to the media. Fluorescence was measured at 633 nm/695 nm excitation/emission. Each data point represents the mean of three independent experiments ± s.d (standard deviation) (**b**) sfGFP expression in the presence or absence of SelB WT or SelB-v2, and in the presence or absence of selenium in the media. Fluorescence was measured at 485 nm/528 nm excitation/emission. Each data point represents the mean of three independent experiments ± s.d (standard deviation).

Sec incorporation in the modified DH10B strain was further confirmed via LC-MS. The sfGFP protein expressed in the presence of selenium and SelB-v2 via a pendant strep-tag, digested with trypsin, and the digested peptides were analyzed for Sec incorporation. Interestingly, while a mass shift was noted, it corresponded to dehydroalanine, rather than selenocysteine (**Supplementary** Fig. 4). It is well-known that Sec can be converted to dehydroalanine through beta-elimination^29^, and thus these results are consistent with selenocysteine incorporation. In order to more directly monitor Sec incorporation, a *E. coli* dihydrofolate reductase (DHFR) gene engineered to contain a selenyl-sulfhydryl bond^30^ was introduced into the modified DH10B strain. Top-down mass spectrometry showed Sec incorporation only upon SelB-v2 induction and selenium addition, and no detectable Ser incorporation (**Fig. 5a**). EThcD fragmentation also confirmed the formation of a selenide-sulfur bond upon incorporation of Sec at position 39 (**Fig. 5b**). In contrast, when identical experiments were performed in C321dA strain and used for expression, roughly 16 % of incorporation of Ser was observed (**Supplementary** Fig. 5), again indicating that this strain may in some cases be problematic for monitoring protein expression and suppression. Previous work has also hinted at this: in the context of the C321dA ‘Amberless’ strain, translation of either glutathione peroxidases(GPx) or mammalian TrxR1 lacking a SECIS element yielded Sec incorporation at the TAG codon but also misincorporation of glutamine and lysine^31^.

**Figure 5.**
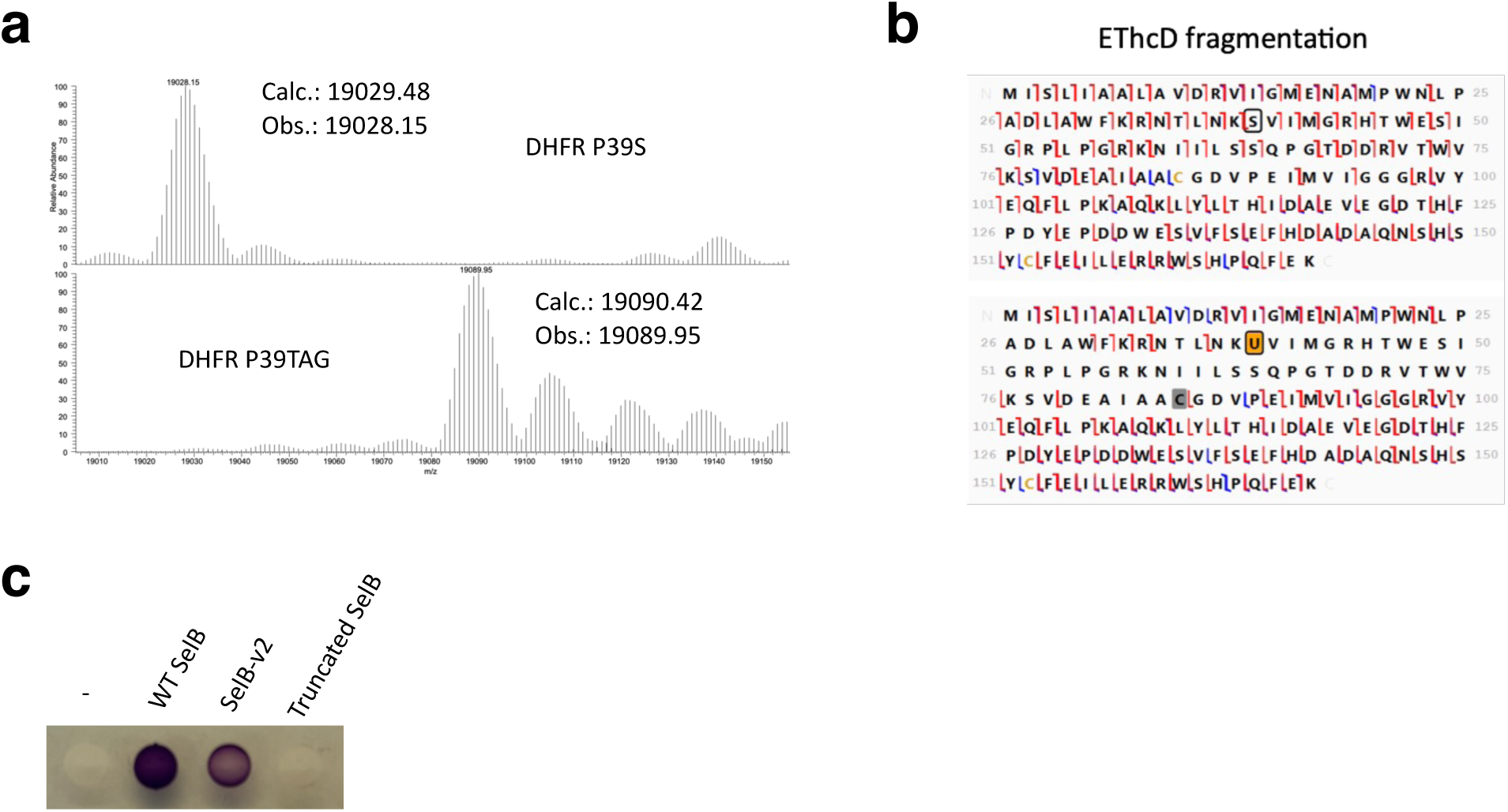
Mass spectrometric and functional characterization of SelB-v2 insertion of selenocysteine. (**a**) SelB-v2 was expressed in C321 dSelABC along with either DHFR P39S (top) or DHFR P39TAG (bottom), and the DHFR proteins were purified in the presence of selenium and MS analyses carried out. The top shows the deconvoluted mass spectrum of DHFR, with serine at position 39 (calculated mass is 19029.48 and observed mass is 19028.15). The bottom is the deconvoluted mass spectrum of DHFR with TAG at position 39 (calculated mass is 19090.42 and observed mass is 19089.95). (**b**) EThcD (Electron-Transfer/higher-energy collision Dissociation) fragmentation pattern of the purified DHFR P39S (top) and DHFR P39TAG (bottom) which shows the selenium-sulfur bond. Slash marks represent the cleavage sites for different N-terminal and C-terminal ions; regions that are ‘protected’ by covalent crosslinking drop out of the mass spectra which was also reported in previously reported selenocysteine containing DHFR^10^. c) Benzyl viologen colorimetric assay. The NMC-A reporter was replaced with a formate dehydrogenase H (fdhF) reporter. The reduction of benzyl viologen (monitored by color change) by expressed fdhF confirms the successful incorporation of selenocysteine into the protein. The functional fdHF expression assay was carried out with SelB-WT, SelB-v2 and SelB- truncated.

Having used engineered reporters to show that SelBv-2 mediated selenocysteine incorporation, we attempted to determine whether SelB-v2 could also incorporate selenocysteine in a natural context, via formate dehydrogenase (fdHF). This native selenoprotein normally has a SECIS element to guid**e** selenocysteine. We removed the genomic fdHF from C321 dSelABC and instead expressed the formate dehydrogenase gene from the same plasmid as SelB-v2. fdHF still relied on its endogenous promoter, while SelB-v2 was expressed from a tac promoter and induced via IPTG. Plate-based assays with benzyl viologen assay showed functional expression of formate dehydrogenase with SelB-v2, although overall enzymatic activity was slightly lower than with the SelB wild-type **(Fig. 5c**).

### Directed evolution of the SelB-v2 into Ser-tRNA^Sec^ specific elongation factor

Since SelB-v2 showed specific recruitment of Sec-tRNA^Sec^ to the ribosome in the absence of a SECIS element, we explored whether it could be used as an orthogonal equivalent of EF-Tu in an even more general manner. Given that the biosynthetic pathway for Sec-tRNA^Sec^ involves the formation of Ser-tRNA^Sec^ as an early intermediate, we were curious whether serine loading might be handled by SelB-v2.

To determine whether specific serine loading could be accomplished, the catalytic serine (position 71) in the beta-lactamase gene was changed to the TAG stop codon, all in the context of the C321dA strain. This made NMC-A a Ser-dependent reporter (NMC-A(Ser)) (**Supplementary** Fig. 6). Upon induction of SelB-v2, the NMC-A S71TAG gene yielded increase in cell growth at low- to mid-concentrations of carbenicillin tested, indicating the serine incorporation was likely taking place (**Fig. 6a**). In order to change or broaden the specificity of SelBv-2 for other amino acids, the residues within the amino acid binding pocket (within 4 Angstroms of the amino acid: Y42, Y44, R181, T193, R236) were randomized, and the SelB-v2 library was plated on LB-agar containing 50 μg/mL Carb. Plate-based assays were used because the TAG codon can mutate to Ser via just one mutation (as opposed to two for Cys), and thus we anticipated (and initially found) that liquid culturing could easily enrich such single mutants. Roughly 150 colonies were observed on the 50 μg/mL Carb plate and 96 colonies were picked and sequenced.

**Figure 6.**
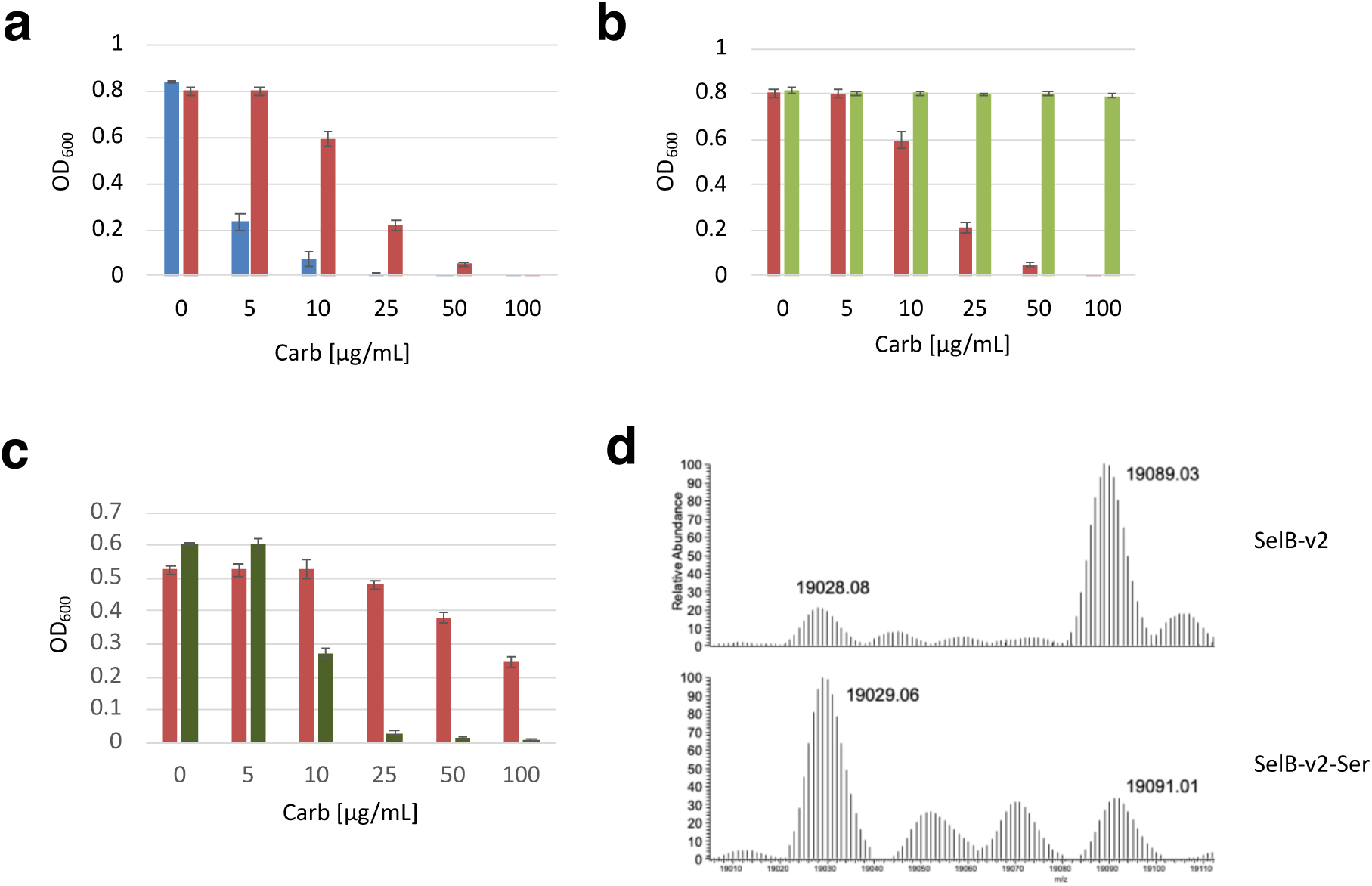
Characterization of SelB mutants for serine incorporation (**a**) Growth assay in the presence of carbenicillin with expression of SelB-v2 and NMC-A S71TAG, which replaces a key serine with a stop codon. The blue bar indicates growth without induction while the red bar is with induction of SelB-v2 by IPTG. Each data point represents the mean of three independent experiments ± s.d (standard deviation). (**b**) Growth assay in the presence of carbenicillin with expression of NMC-A S71TAG and either induced SelB-v2 (red) or the evolved SelB-v2-Ser (green). Each data point represents the mean of three independent experiments ± s.d (standard deviation). (**c**) Growth assay in the presence of carbenicillin with expression of NMC-A C70TAG (which is sensitive to selenocysteine insertion) and either induced SelB-v2 (red) or SelB-v2-Ser (green). Each data point represents the mean of three independent experiments ± s.d (standard deviation). (d) Intact mass spectra of DHFR P39TAG expressed and purified in the presence of selenium and SelB-v2 (top) or SelB-v2-Ser (bottom). The calculated mass of DHFR P39S is 19029.48 and DHFR P39Sec with wild-type SelB is 19090.42, whereas when SelB- v2 is co-expressed, the observed mass for DHFR P39S is 19028.08 and for DHFR P39Sec is 19089.03. When SelB-v2-Ser is co-expressed, the observed mass for P39S is 19029.06 and for DHFR P39Sec is 19091.01.

Sequencing of over 96 colonies picked from the plate revealed convergence of mutations with the top mutant constituting (SelB-v2-Ser; Y42Y, Y44F, R181R, T193C, R236S) representing a quarter of the clones. SelB-v2-Ser showed improved incorporation of Ser compared to the original SelB-v2 (**Fig. 6b**), as active as the wild-type enzyme.

In order to see if the mutated binding pocket of SelB-v2-Ser also affects the incorporation of Sec, the original NMC-A C70TAG gene, which was used to engineer SelB-v2, was used to assess the Sec incorporation efficiency. It should be noted that the C321 strain retains SelD in the genome, which means that Ser-tRNA^Sec^ can be converted to Sec-tRNA^Sec^ by SelA using selenophosphate as the substrate when selenium is present. While SelB-v2 previously allowed growth at higher concentrations of carbenicillin (25-100 μg/mL), this growth advantage was lost in SelB-v2-Ser (**Fig. 6c**). Ser incorporation was further measured in DHFR P61TAG in the presence of selenium and induction of SelB-v2-Ser. Intact mass analysis revealed that despite selenocysteine being fully available for incorporation, roughly 74% of the protein contained serine, as opposed to only 17% with the parental SelB-v2 (**Fig. 6d**) indicating that tRNA selectivity can be modulated by mutating the amino acid binding pocket. The change in aminoacyl-tRNA selectivity was also confirmed via SDS-PAGE gel analysis (**Supplementary** Fig. 7).

## Discussion

The traditional view of the role of SECIS element is to enhance selenocysteine incorporation by recruiting SelB to the ribosome, enabling tRNA^Sec^ loading. This view has been validated by multiple results in which mutation of the SECIS element decreased selenoprotein translation^32–35^. The notion that the SECIS enhances selenoprotein production is also seemingly inherent in its mechanism, wherein tRNA^Sec^ is first recognized by SelA and then transferred to SelB:GTP complex, which in turn is recognized by the SECIS element, leading to correct delivery of Sec-tRNA^Sec^ to ribosome^36^.

However, a consideration of the possible origins of selenocysteine incorporation tell a slightly different story. If selenoprotein production proved important for the evolution of metabolism, the simplest innovation would have been to have EF-Tu adopt a selenocysteine-charged tRNA; the introduction of the complex system that currently exists for selenocysteine incorporation would have emerged much later. The ostensible reason for its emergence would likely have been to not enhance selenocysteine incorporation, but rather to restrict it, limiting it to only certain stop codons. This explanation has also been embraced in the literature^37,38^.

Over time, SelB could have emerged by duplication and divergence of EF-Tu^39,40^, or possibly other translation factors such as IF-2^41,42^. This would have been the key step that allowed further elaboration of the remaining selenocysteine incorporation machinery, and would have occurred prior to the Last Universal Common Ancestor (LUCA)^39^. What is less clear is why this would have been necessary. Other suppressor systems arise with regularity, and do not greatly diminish organismal fitness^43,44^.

Previous work from our lab on a different system for selenocysteine incorporation, one that involved altering Sec-tRNA to be used by EF-Tu for loading^10,25^, showed that large scale selenocysteine incorporation, even in an Amberless background, was highly toxic to cells. Thus, there would likely have been significant selective pressure following the initial, EF-Tu mediated incorporation of selenocysteine into proteins to generate a system that would limit selenocysteine introduction to only specified codons.

Overall, these previous results suggest that the role of SelB is not to promote efficiency, but rather to greatly limit suppression. In this model, coupling of SelB to the SECIS is not functionally required, but rather is an augmented function, following the original divergence of SelB. The largely structurally separate RNA-binding domain on SelB is unrelated to other translation factors, and would have most likely been added following divergence, further supporting this hypothesis; the more ancestral version of SelB is more akin to our own SelB-v2.

If this hypothesis is valid, then not only is the RNA-binding domain of SelB potentially a late addition, it should also be functionally separable, without compromising SelB’s tRNA loading activity. This hypothesis was the basis for the experiments described herein, where we in fact demonstrate that a SelB variant that was completely separable from the SECIS could readily be generated by only a few rounds of directed evolution. Interestingly, when the SECIS-binding domain is left in place, the S206P and P277S mutations (singly or in tandem) are predicted to lead to a complete reorientation of the SECIS-binding domain relative to the rest of the protein (**Supplementary** Fig. 8), potentially perturbing function. This further suggests the likely ancestral independence of the ancestral-like SelB-v2.

This hypothesis also impacts further alteration of the genetic code. While the SECIS may have once been required to stabilize an expanded code, it obviously now limits expansion, and the more ancestral-like SelB-v2 provides an intriguing opportunity for genetic code expansion. EF-Tu is of course known to balance the binding of various tRNA-amino acid conjugates, in order to help ensure the overall accurate loading across from a wide variety of codons. This ‘balancing act’ is largely mediated through a complex series of interactions between tRNAs and the surface of SelB, and likely has its limits. The introduction of a new, orthogonal loading factor, SelB-v2, can potentially open the way to expansions of the genetic code much larger than just 21 amino acids, as it takes over the loading of an entirely new expansion set via its own balancing act. The fact that we were able to not only introduce selenocysteine, but also serine via SelB-v2 directed evolution provides an existence proof for further expansion of its role in loading non-canonical amino acids in further expanded codes.

## Material and methods

### Strain Construction

Strains used in this study were kindly gifted from Prof. Ross Thyer and were used in explorations of selenocysteine incorporation.^10^ One exception was the strain used in the benzyl viologen assay, wherein the fdHF gene was further deleted from the genome of C321 lacking SelAB and SelC using the lamda Red system from Datsenko and Wanner^45^, and replaced with a chloramphenicol resistance marker.

### Library Construction and Selection

The overall design of the plasmid (pSelAC-araBAD-SelB) for selecting SECIS-independent SelB is shown in **Supplementary Table 3**. Libraries were created by using error-prone PCR to amplify the coding region of SelB (GeneMorph II Random Mutagenesis Kit, Agilent, #200550). The library (pSelAC-araBAD-SelB) was transformed into an electrocompetent C321 dSelABC strain which was then grown for 1 hour at 37℃ in 2 mL of SOC, and later transferred to 200 mL of LB medium in the presence of 10 μM Na_2_SeO_3_, 0.01 % (v/v) arabinose, 50 μg/mL kanamycin and various concentrations of carbenicillin (**Supplementary Table 1**) for 16 hours at 37℃. Cells were recovered, plasmids isolated, and the coding region of SelB was amplified by PCR (either using Platinum SuperFi II DNA polymerase (Thermo Fisher, #12361010) or the GeneMorph II Random Mutagenesis Kit). The PCR products (Oligo1-4) were introduced into the same plasmid via Gibson assembly. The same procedure was used to derive SelB-v2, except cloning was carried out via Golden Gate assembly (Oligo5-8).

The overall design of the plasmid (pSelC-tat-SelB) for identifying serine-incorporating variants of SelB-v2 is shown in **Supplementary Table 4** and **Supplementary** Fig. 1. The plasmid pSelC-tat- SelB was initially prepared by amplifying four fragments using four sets of primers that had NNS primers targeting 5 positions (Y42, Y44, R181, T193, R236) (Oligo 9-11, 12-13, 14-15, 6-16; **Supplementary Table 6**). These four fragments were combined using assembly PCR and then cloned into the vector (pSelC-tat-SelB) using Golden Gate assembly (Oligo 6,8,10,17; **Supplementary Table 6**). The library was transformed into electrocompetent C321 dSelABC, incubated for 1 hour at 37℃ in 2.4 ml of SOC and then plated onto four 15 cm LB-plates containing 50 μM IPTG, 50 μg/mL carbenicillin and 50 μg/mL kanamycin. Roughly 150 colonies were picked after 16 hours at 37℃ and then individually checked for sequencing.

### NMC-A C70TAG assay

Plasmids (pSelAC-araBAD-SelB) encoding SelB variants were freshly transformed into C321dSelABC and grown on LB-agar plates containing 50 μg/mL kanamycin. Three colonies were picked from a plate and grown overnight in LB media containing 50 μg/mL kanamycin at 37 °C. The overnight culture was diluted 1:100 in fresh LB supplemented with 50 µg/mL kanamycin, 10 μM Na_2_SeO_3_ and 0.01 % (v/v) arabinose, and 0, 5, 10, 25, 50, 100 µg/mL carbenicillin. Cultures were incubated for 16 h at 37 °C with shaking. OD600 values were measured using a TECAN plate reader.

### NMC-A S71TAG assay

Some 10 ng of plasmids (pSelC-tac-SelB) containing corresponding SelB variants were transformed into electrocompetent C321dSelABC in triplicate, and grown for 2 hours. Cultures were then diluted 1:100 in fresh LB supplemented with 50 µg/mL kanamycin, 50 µM IPTG and 0, 5, 10, 25, 50, 100 µg/mL carbenicillin, and incubated for 16 h at 37 °C with shaking. OD600 values were measured using a TECAN plate reader.

### sfGFP and smURFP assay

The sfGPF reporter plasmid was prepared by swapping the NMC-A gene in pSelAC-araBAD-SelB with sfGFP, and the promoter was changed to the Ipp5 promoter. The smURFP reporter plasmid was prepared by swapping the NMC-A gene in pSelAC-araBAD-SelB with the smURFP-HO-1 gene (retaining the NMC-A promoter). Plasmids encoding the WT-SelB or SelB-v2 were freshly transformed into DH10B dSelABC and grown on LB-agar plates containing 50 μg/mL kanamycin. Three colonies were picked from a plate and grown overnight in LB media containing 50 μg/mL kanamycin at 37 °C. Overnight cultures were diluted 1:100 in fresh LB supplemented with 50 µg/mL kanamycin, 10 μM Na_2_SeO_3_ and 0.01 % (v/v) arabinose, and incubated for 16 h at 37 °C with shaking. OD600 as well as fluorescence (ex 485 nm /em 528 nm for sfGFP, and ex 633 nm/em 695 nm for smURFP) was measured using a TECAN plate reader.

### Protein purification

To express and purify protein for MS analyses, the NMC-A gene in pSelAC-araBAD-SelB was replaced with the folA gene containing the mutation P61S or P61TAG followed by a strep-tag II (WSHPQFEK). Plasmids encoding SelB-v2 or SelB-v2-Ser were freshly transformed into C321 dSelABC or DH10B dSelABC and grown on LB-agar plates containing 50 μg/mL kanamycin. A single colony was picked from a plate and grown overnight in LB media containing 50 μg/mL kanamycin at 37 °C. Overnight cultures were diluted 1:100 in fresh LB (50 ml) supplemented with 50 µg/mL kanamycin, 1 μM Na_2_SeO_3_ and 0.002 % (v/v) arabinose for the DH10B dSelABC strain, and 10 μM Na_2_SeO_3_ and 0.01 % (v/v) arabinose for the C321 dSelABC strain, and then grown overnight. Cells were harvested by centrifugation at 10,000 x g for 10 min and resuspended in 3 mL of BugBuster (Novagen #70584-3). Following 20 min incubation on a rotating mixer at 20 rpm at room temperature, cells were clarified by centrifugation at 16,000 x g for 20 min at 4 C. Lysates were passed through a 0.2 μm filter and DHFR or sfGFP samples were purified using MagStrept Strep-Tactin XT beads following the manufacturer’s instructions (IBA Lifesciences #2-4090-002). After purification, the buffer was exchanged to 0.1 % formic acid in water by acetone precipitation.

### Benzyl Viologen Assay

To carry out the benzyl viologen assay, a reporter plasmid was prepared by replacing the promoter and NMC-A gene with the endogenous promoter of the fdhF gene and a fdhF gene with a TGA140TAG mutation. *E. coli* C321 lacking the SelAB, SelC, and fdhF loci was transformed with the reporter plasmid that contained either wild-type SelB, a SelB truncation, or SelB-v2. Transformants were grown overnight at 37 °C in LB medium containing 50 μg/mL kanamycin. Overnight cultures were diluted 1:20 and incubated for three hours in 1.5 mL of LB medium and cultures were then normalized to OD_600_ = 0.5 by the addition of LB medium. A number of 5 μl aliquots were dotted on LB agar plates containing 50 μg/mL kanamycin, 5 mM sodium formate, 1 μM Na_2_MoO_4_, 1 μM Na_2_SeO_3_. Plates were incubated at 37 °C for 3 h under aerobic conditions and then grown for 60 h at 37 °C under anaerobic conditions using the pouch system (BD B260683). Upon removal from the pouch, plates were immediately overlaid with 0.75 % agar containing 1 mg/mL benzyl viologen, 250 mM sodium formate and 25 mM KH_2_PO_4_ at pH 7.0. Plates were then photographed within 1 h of overlaying.

### AlphaFold3 Structure Prediction

AlphaFold3 (https://alphafoldserver.com/) was used to generate the following structures for non-truncated SelB: wild type, S206P, P277S, and [S206P, P277S]^46^. The following seeds were used for each of these structures: 865694845, 601147261, 1701205798, and 1572407154, respectively, and the top ranked structures “_model_0” were used.

## Acknowledgements

The research performed by S.I. was sponsored by grants from the Toyobo Biotechnology Foundation and JSPS Overseas Research Fellowships. S.I. and A.G. were also supported by the Army Research Laboratory under Cooperative Agreement Number W911NF-22-2-0246 and by the Air Force Research Laboratory under S-168-1X6-001. The views and conclusions contained in this document are those of the authors and should not be interpreted as representing the official policies, either expressed or implied, of the Army Research Laboratory, Air Force Research Laboratory, or the U.S. Government. The U.S. Government is authorized to reproduce and distribute reprints for Government purposes notwithstanding any copyright notation herein. The contributions of C.W.K. and A.D.E. were supported by the Welch Foundation (F- 1654). Protein identification was provided by the UT Austin Center for Biomedical Research Support Biological Mass Spectrometry Facility (RRID: SCR_021728). We thank Prof. Ross Thyer for kindly gifting the strains and plasmid.

## Author Contributions

S.I. and A.D.E. conceived the project idea and oversaw all aspects of the research. S.I. and A.G. conducted biochemical studies. C. W. K. did the computational modeling. All authors contributed to writing the manuscript.

## Data Availability

All raw data are available from the corresponding author upon reasonable request.

