## Supplementary information for "Directed evolution of a SelB variant that does not require a SECIS element for function"

**Supplementary Table 1| Selection conditions of each round**

(a) To obtain SelB-v1, 4 rounds of selection were carried out. (b) To obtain SelB-v2, 7 rounds of selection were carried out, in liquid culture. N/A, not applicable.

**a**

| round | Carbenicillin concentration [μg/mL] | Error prone PCR after screening | Screening method |
|---|---|---|---|
| 1st | 5 | No | Liquid culture |
| 2nd | 10 | Yes | Liquid culture |
| 3rd | 10 | No | Liquid culture |
| 4th | 10 | No | Liquid culture |

**b**

| round | Carbenicillin concentration [μg/mL] | Error prone PCR after screening |
|---|---|---|
| 2-1st | 10 | No |
| 2-2nd | 10 | No |
| 2-3rd | 10 | Yes |
| 2-4th | 10 | No |
| 2-5th | 10 | Yes |
| 2-6th | 10 | No |
| 2-7th | 100 | N/A |

**Supplementary Table 2| List of mutations found in clones picked after round 5**

A total of six clones were picked and compared with wild-type SelB.

| Position | Original codon (amino acid) | Mutated codon (amino acid) | Frequency |
|---|---|---|---|
| 38 | ATC (I) | ATA (I) | 6/6 |
| 83 | GCG (A) | GCA (A) | 6/6 |
| 98 | ATT (I) | GTT (V) | 6/6 |
| 166 | GAA (E) | GAT (D) | 6/6 |
| 189 | GGG (G) | GGC (G) | 6/6 |
| 206 | TCA (S) | CCA (P) | 6/6 |

| Position | Original codon (amino acid) | Mutated codon (amino acid) | Frequency |
|---|---|---|---|
| 277 | CCG (P) | TCG (S) | 6/6 |
| 362 | GCG (A) | GCA (A) | 6/6 |
| 413 | GCG (A) | ACG (T) | 5/6 |
| 473 | AGC (S) | AAC (N) | 6/6 |
| 477 | TGG (W) | CGG (R) | 6/6 |
| 487 | TTC (F) | CTC (L) | 6/6 |

### Supplementary Table 3| Plasmid sequence of pSelAC-araBAD-SelB

The ORF of NMC-A is pink, with the first TAG underlined and the second TAG codon bolded; these codons were both originally TGT and TGC, respectively. The ORF of SelB is green, the ORF of SelA is yellow, and SelC is cyan, with the anticodon underlined.

TCGACCGATGCCCTTGAGACGTTGATCGGCACGTATGGCAGATAGTAAATTTTATAGATTTAAATCAACATCTATAACTGTATAATTCGTTTTCTCAACTCATTACAACACTCCGTTAGTAATGAAGCT
CATTCTATACAATGACAGTTAATAGGTAAAGTTATGTCACTTAATGTAAAGCAAAGTAGAATAGCCATCTTGTTTAGCTCTTGTTTAATTTCAATATCATTTTTCTCACAGGCCAATACGAAGGGCATTGA
TGAGATTAAAAACCTTGAAACAGATTTCAATGGCAGGATTGGTGTCTACGCTTTAGACACTGGCTCGGGTAAATCATTTTCGTACAGAGCAAATGAACGATTTCCATTATAGAGTTCTTTTAAAGGTTT
TTTAGCTGCTGCTGTATTAAAAGGCTCTCAAGATAATCGACTTAATCTTAATCAGATTGTGAATTATAATACAAGAAGTTTAGAGTTCCATTCACCCATCACAACTAAATATAAAGATAATGGAATGTCAT
TAGGTGATATGGCTGCTGCTGCTTTACAATATAGCGACAATGGTGCTACTAATATTATTCTTGAACGTTATATCGGTGGTCCAGAGGGTATGACTAAATTCATGCGGTCGATTGGAGATGAAGATTTTA
GACTCGATCGTTGGGAGTTAGATCTAAACACAGCTATTCCAGGCGATGAGCGTGACACATCTACACCTGCAGCAGTAGCCAAGAGTCTGAAAACCCTTGCTCTGGGTAACATACTTAGTGAACAT
GAAAAGGAAACCTATCAGACATGGTTAAAGGGTAACACAACCGGTGCAGCGCGTATTCGTGCTAGCGTACCAAGCGATTGGGTAGTTGGCGATAAAACTGGTAGT**TAG**GGAGCATACGGTACG
GCAAATGATTATGCGGTAGTCTGGCCAAAGAACCGGGCTCCTCTTATAATTTCTGTATACACAACAAAAAACGAAAAAGAAGCCAAGCATGAGGATAAAGTAATCGCAGAAGCTTCAAGAATTGCA
ATTGATAACCTTAAATAACAAAGCCCGCCGAAAGGCGGGCTTTTTTTTGGATCCTGTAGAAACGCAAAAAGGCCATCCGCATGACTAACATGAGAATTACAACTATCACCGCAAGGGATAAATATC
TAACACCGTTATGACAACTTGACGGCTACATCATTCACTTTTTCTTCACAACCGGCACGGAACTCGCTCGGGCTGGCCCCGGTGCATTTTTTAAATACCCGCGAGAAGTAGAGTTGATCGTCAAA
ACCAACATTGCGACCGACGGTGGCGATAGGCATCCGGGTGGTGCTCAAAAGCAGCTTCGCCTGGCTGATACGTTGGTCCTCGCGCCAGCTTAAGACGCTAATCCCTAACTGCTGGCGGAAAA
GATGTGACAGACGCGACGGCGACAAGCAAACATGCTGTGCGACGCTGGCGATATCAAAATTGCTGTCTGCCAGGTGATCGCTGATGTACTGACAAGCCTCGCGTACCCGATTATCCATCGGTG
GATGGAGCGACTCGTTAATCGCTTCCATGTGCCGCAGTAACAATTGCTCAAGCAGATTTATCGCCAGCAGCTCCGAATAGCGCCCTTCCCCTTGCCCGGCGTTAATGATTTGCCCAAACAGGTC
GCTGAAATGCGGCTGGTGCGCTTCATCCGGGCGAAAGAACCCCGTATTGGCAAATATTGACGGCCAGTTAAGCCATTCATGCCAGTAGGCGCGCGGACGAAAGTAAACCCACTGGTGATACC
ATTCGCGAGCCTCCGGATGACGACCGTAGTGATGAATCTCTCCTGGCGGGAACAGCAAAATATCACCCGGTCGGCAAACAAATTCTCGTCCCTGATTTTTCACCACCCCCTGACCGCGAATGG
TGAGATTGAGAATATAACCTTTCATTCCCAGCGGTCGGTCGATAAAAAAATCGAGATAACCGTTGGCCTCAATCGGCGTTAAACCCGCCACCAGATGGGCATTAAACGAGTATCCCGGCAGCAG
GGGATCATTTTGCGCTTCAGCCATACTTTTCATACTCCCGCCATTCAGAGAAGAAACCAATTGTCCATATTGCATCAGACATTGCCGTCACTGCGTCTTTTACTGGCTCTTCTCGCTAACCAAACCG
GTAACCCCGCTTATTAAAAGCATTCTGTAACAAAGCGGGACCAAAGCCATGACAAAAACGCGTAACAAAAGTGTCTATAATCACGGCAGAAAAGTCCACATTGATTATTTGCACGGCGTCACACTT
TGCTATGCCATAGCATTTTTATCCATAAGATTAGCGGATCCTACCTGACGCTTTTTATCGCAACTCTCTACTGTTTCTCCATACCCGTTAGGAGGATTGTATGATTATCGCGACTGCCGGACATGTGG
ATCATGGAAAGACAACATTGTTGCAGGCGATTACTGGCGTAAATGCTGACCGTCTGCCGGAAGAAAAAAAGCGCGGCATGACCATCGATCTCGGCTATGCCTACTGGCCGCAGCCGGATGGT
CGCGTGCCTGGTTTTATCGACGTTCCCGGTCATGAAAAGTTTCTTTCCAACATGCTGGCGGGCGTTGGTGGTATCGATCACGCGCTGTTGGTGGTGGCGTGCGATGACGGCGTGATGGCACAG
ACCCGTGAGCATCTGGCGATTTTGCAGCTGACCGGTAACCCGATGCTGACAGTGGCGCTGACCAAAGCCGATCGCGTGGACGAAGCGCGTGTTGATGAGGTTGAACGCCAGGTAAAGGAGG
TTCTGCGGGAATACGGTTTTGCTGAGGCAAAACTGTTTATCACCGCAGCAACCGAAGGTCGGGGAATGGATGCCCTGCGCGAGCATCTGCTTCAGTTGCCGGAACGCGAGCACGCCAGCCAA
CATAGTTTCCGCCTCGCGATTGACCGCGCATTTACCGTAAAAGGTGCCGGGCTGGTCGTCACCGGTACGGCGTTAAGCGGGGAAGTGAAGGTAGGCGATTCACTCTGGCTGACTGGTGTAAAT
AAACCGATGCGTGTACGTGCGCTGCATGCGCAAAACCAGCCAACAGAAACCGCCAATGCCGGGCAGCGTATCGCGCTTAACATCGCGGGTGATGCGGAAAAAGAGCAGATTAACCGTGGC
GACTGGCTGCTTGCCGATGTGCCGCCAGAGCCGTTCACACGGGTGATTGTCGAGCTTCAAACCCATACACCGCTGACCCAGTGGCAGCCGCTGCATATTCACCACGCCGCCAGCCACGTC
ACGGGACGCGTTTCACTGCTGGAAGATAACCTTGCTGAACTGGTCTTCGACACCCCGTTATGGCTGGCAGATAACGACCGCCTGGTATTGCGCGATATCTCTGCCCGCAACACGCTGGCCGG
AGCGCGCGTCGTGATGCTTAACCCGCCGCGTCGCGGTAAACGTAAGCCGGAATATCTGCAATGGCTGGCGTCACTTGCACGGGCGCAGAGCGATGCCGATGCGTTATCTGTTCATCTGGAAC
GCGGCGCGGTTAACCTTGCGGATTTCGCCTGGGCGCGCCAGCTCAACGGCGAAGGGATGCGCGAATTGCTGCAACAGCCTGGTTATATTCAGGCTGGTTATAGCTTGTTGAATGCGCCGGTT
GCCGCCCGCTGGCAGCGGAAAATTCTCGACACATTAGCGACTTATCATGAGCAACATCGCGATGAACCTGGCCCTGGGCGCGAACGTCTGCGACGTATGGCGTTGCCAATGGAAGATGAAG
CGCTGGTACTGTTGCTGATTGAAAAGATGCGCGAAAGCGGCGACATCCACAGCCATCACGGCTGGCTGCATCTGCCAGATCACAAAGCGGGCTTCTAACATTGTCTTATAAATACCAAAAATTTC
ACGTCTCTTGTTTAAAGAGGCGTGCTTTAATATTTTCTTAGAGTCCAATACATGAGAGTTTTAATTTCGTTTTTTTTCCTTATTTTTTCTATTAGCGTTAGAGCTGACTGTGGCCTTGATAGATCTTCGAAGC
TTGGGCCCGAACAAAAACTCATCTCAGAAGAGGAGAGATAAATTGCACTGAAATCTAGAGTAACGGAATAGCTGTTCGTTGACTTGATAGACCGATTGATTCATCATCTCATAAATAAAGAAAAACC
ACCGCTACCAACGGTGGTTTTCTCAAGGTTCGCTGAGCTACCAACTCTTTGAACCAAGGTAAGTGGGTTGGAGGACCGCACTCACCAAAATCTGTTCTTTCAGTTTAGCCTTAACAGGTGCATAA
CTTCAAGACAAAGTCCTCTAAATCAGTTACCAATGGCTGCTGCCAGTGGCGATAAGTCGTGTCTTACCGGGTTGGACTCAAGACGATAGTTACCGGATAAGGCGCAGCGGTCGGGCTGAACGG
GGGGTTCGTGCACACAGCCCAGCTTGGAGCGAACGACCTACACCGAACTGAGATACCAACAGCGTGAGCTATGAGAAAGCGCCACGCTTCCCGAAGGGAGAAAGGCGGACAGGTATCCGG
TAAGCGGCAGGGTCGGAACAGGAGAGCGCACGAGGGAGCTTCCAGGGGGAAACGCCTGGTATCTTTATAGTCCTGTCGGGTTTCGCCACCTCTGGCTTGAGCGTCGATTTTTGTGATGCTCGT
CAGGGGGGCGGAGCCTATGGAAAAACGCCTGCGGCGTTGGCTTCTTCCGGTGCTTTGCTTTTTGCTCACATGTTCTTTCCGGCTTTATCCCCTGATTCTGTGGATAACCGTATTACCGCTTTTGAG
TGAGCTGACACCGCTCGCCGCAGTCGAACGACCGAGCGTAGCGAGTCAGTGAGCGAGGAAGCGGAAGAGCGCTGCATGCCTATTTGTTTATTTTTCTAAATACATTCAAATATGTATCCGCTCA
TGAGACAATAACCCTGATAAATGCTTCAATAATATTGAAAAAGGAAGAGTATGAGCCATATTCAACGGGAAACGTCTTGCTCTAGGCCGCGATTAAATTCCAACATGGATGCTGATTTATATGGGTAT
AAATGGGCTCGCGATAATGTCGGGCAATCAGGTGCGACAATCTATCGATTGTATGGGAAGCCCGATGCGCCAGAGTTGTTTCTGAAACATGGCAAAGGTAGCGTTGCCAATGATGTTACAGATG
AGATGGTCAGACTAAACTGGCTGACGGAATTTATGCCTCTTCCGACCATCAAGCATTTTATCCGTACTCCTGATGATGCATGGTTACTCACCACTGCGATCCCCGGGAAAACAGCATTCCAGGTA
TTAGAAGAATATCCTGATTCAGGTGAAAATATTGTTGATGCGCTGGCAGTGTTCCTGCGCCGGTTGCATTCGATTCCTGTTTGTAATTGTCCTTTTAACAGCGACCGCGTATTTCGTCTCGCTCAGG
CGCAATCACGAATGAATAACGGTTTGGTTGATGCGAGTGATTTTGATGACGAGCGTAATGGCTGGCCTGTTGAACAAGTCTGGAAAGAAATGCATAAACTTTTGCCATTCTCACCGGATTCAGTCG
TCACTCATGGTGATTTCTCACTTGATAACCTTATTTTTGACGAGGGGAAATTAATAGGTTGTATTGATGTTGGACGAGTCGGAATCGCAGACCGATACCAGGATCTTGCCATCCTATGGAACTGCCT
CGGTGAGTTTTCTCCTTCATTACAGAAACGGCTTTTTCAAAAATATGGTATTGATAATCCTGATATGAATAAATTGCAGTTTCATTTGATGCTCGATGAGTTTTTCTAACTGCAATAAGGTTGTTTTGCCGT
GGTCAACGTGTCCGGCAGTCGCAATAATCATTTCAACAACATCTCCAAAAACCGTTGCTCATCTTCAAGGCAGCGTAAATCCAGCCACAATCGTCCGTCATAAATACGACCAATCACCGGCACT
GGCAATTCACGCCAGCGGGCGGCTAATGACTCAAGGTGGCTACCGCGTCCATCATGGGGTGTAAACGTTAATGCCGCGCTCGGCAGGCGATCAACCGGCAGCGAACCACTGCCAATCTGC
GAAAGACATGGCATAACCTGTACCGCAAACTCCGCGCCGTAATGTGCGGCAAGGGGGGCCTGTAAACGTTGTGCCTGGATTTGAATGACCTCTGCGCTGCGGGTAAGCAGGCGCAGGGTCG
GTAATTTTTCACTCAGAGCTTCAGGGTGTAAATAAAGACGCAACGTGGCTTCCAGCGCCGCGAGGGTCATTTTATCCGCGCGTAATGCACGCTTCAGCGGGTGGCTTTGCAGGCGGGCGATCA
TCTCTTTTTTACCAACAATAATTCCTGCCTGCGGCCCGCCTAACAACTTGTCGCCGGAGAAACTCACCAGACTGACGCCCGCCGCAATCAACTCCTGCGGCATTGGCTCTTTCGGCAAACCGT
ACTGGCTAAGATCGACCAGCGAGCCACTGCCTAAATCAGTCACTACGGGAACATCCAGCTCTTTGCCGAGCGCCACCAGTTCCGCTTCATCTATCGCTTTGGTGAACCCCTGAATGCTGTAGTT
ACTGGTATGTACTTTCATCAACAGTGCGGTATTTTCATTCACCGCCTGACGATAATCATTCGCGTGCGTGCGGTTGGTGGTCCCTACTTCGTGTAGGGTGCAGCCTGCCTGACGCATAACATCGG
GAATACGAAACGCGCCGCCAATCTCCACCAGTTCGCCGCGAGATACCACCACCTCTTTTCCGCTGGCAGTGGCCGCCAACATCAATAACACCGCCGCCGCATTGTTATTGACGATACAGGCA
TCTTCCGCCCCCGTAATACGGCACAGCAGCTGCGCCAGCGCCCGATCGCGATGTCCGCGTCCGGCGTCGTCCAGATCATACTCGAGGGTCACTGGCGAACGCATAGCCTGCGCAACGGC
TTCCACCGCGGCTTCCGCCTGTAAAGCTCGCCCAAGGTTGGTATGCAGCACGGTTCCCGTCAGGTTGATCACCGGACGCAGCGCGCTCTGCGCTTCTTTCGTCAACCGGGCATCGACTTCTT
GCGCCCAGTTTTCACACCACGCAGGCAGCGTCTGGCTGCCACGAATCACTTCTCGCGCTTCGTCGAGCATCTGACGCAACAATTCCACCACGCGGGTGTGACCATAAGTATCACGCAAAGAA
AGGAAGGAGCTATCGCGCAATAAGCGATCAATAGCCGGAAGTTGACTATAGAGGAAACGCGTTTCGGTTGTCATGGTTTAGTTCCTCACCTTGTCGTATTATACTATGCCGATATACTATGCCGAT
GATTAGGAAACCTGGCTGATCAAGGCCCTCTCACACGGAGAAGGGCGTTTAACATAACCACGGATTGTAACGTGAGATGGGTCAGGAGGACATATCGCGCATCAAGCCTTTGGCGGTTCGGTG
CATCCCATGGCGATTGAATATTGAGCAGTAGAAAATATCTGGATTGACAGTATTACAAGGGGTTGGAGAACGTCAGGAGAGGCATTTTGGCGGAAGATCACAGGAGTCGAACCTGCCCGGGACC
GCTGGCGGCCCCAACTGGATTTAGAGTCCAGCCGCCTCACCGGAGACGACGATCTTCCGCGCCTCGATTGCTACATGGAGGCGGGGCGCATTATAGCTACTTCCTTGAGTTTCTACATCCCC
CAGATCAGTTTTTCCGCCTAATACACGCAGCGTATTATCACGAAAGCATTATAGTTCATATAAAACCCATATTTACAATGATATATGAGCGGAATTAACGCTACCACGATCACATAAAACAAGAATC
ATAAATTAATAACCAGATATCGGAATATTCGCTCTCACAGGGATGGCGGCCGC

### Supplementary Table 4| Plasmid sequence of pSelC-tac-SelB

The ORF of NMC-A is pink, with the TAG bolded and underlined (this was originally a serine codon). The ORF of SelB is green, the ORF of LacI is yellow, and SelC is cyan, with the anticodon underlined.

TCGACCGATGCCCTTGAGACGTTGATCGGCACGTATGGCAGATAGTAAATTTTATAGATTTAAATCAACATCTATAACTGTATAATTCGTTTTCTCAACTCATTACAACACTCCGTTAGTAATGAA
GCTCATTCTATACAATGACAGTTAATAGGTAAAGTTATGTCACTTAATGTAAAGCAAAGTAGAATAGCCATCTTGTTTAGCTCTTGTTTAATTTCAATATCATTTTTCTCACAGGCCAATACGAAGG
GCATTGATGAGATTAAAAACCTTGAAACAGATTTCAATGGCAGGATTGGTGTCTACGCTTTAGACACTGGCTCGGGTAAATCATTTTCGTACAGAGCAAATGAACGATTTCCATTATGT**TAG**TC
TTTTAAAGGTTTTTTAGCTGCTGCTGTATTAAAAGGCTCTCAAGATAATCGACTTAATCTTAATCAGATTGTGAATTATAATACAAGAAGTTTAGAGTTCCATTCACCCATCACAACTAAATATAAA
GATAATGGAATGTCATTAGGTGATATGGCTGCTGCTGCTTTACAATATAGCGACAATGGTGCTACTAATATTATTCTTGAACGTTATATCGGTGGTCCAGAGGGTATGACTAAATTCATGCGGT
CGATTGGAGATGAAGATTTTAGACTCGATCGTTGGGAGTTAGATCTAAACACAGCTATTCCAGGCGATGAGCGTGACACATCTACACCTGCAGCAGTAGCCAAGAGTCTGAAAACCCTTGC
TCTGGGTAACATACTTAGTGAACATGAAAAGGAAACCTATCAGACATGGTTAAAGGGTAACACAACCGGTGCAGCGCGTATTCGTGCTAGCGTACCAAGCGATTGGGTAGTTGGCGATAAA
ACTGGTAGTTGCGGAGCATACGGTACGGCAAATGATTATGCGGTAGTCTGGCCAAAGAACCGGGCTCCTCTTATAATTTCTGTATACACAACAAAAAACGAAAAAGAAGCCAAGCATGAGG
ATAAAGTAATCGCAGAAGCTTCAAGAATTGCAATTGATAACCTTAAATAACAAAGCCCGCCGAAAGGCGGGCTTTTTTTTGGATCCTGTAGAAACGCAAAAAGGCCATCCGCATGACTAACA
TGAGAATTACAACTGCGGCGCGCCATCGAATGGCGCAAAACCTTTCGCGGTATGGCATGATAGCGCCCGGAAGAGAGTCAATTCAGGGTGGTGAATATGAAACCAGTAACGTTATACGAT
GTCGCAGAGTATGCCGGTGTCTCTTATATGACCGTTTCCCGCGTGGTGAACCAGGCCAGCCACGTTTCTGCGAAAACGCGGGAAAAAGTGGAAGCGGCGATGGTGGAGCTGAATTACAT
TCCCAACCGCGTGGCACAACAACTGGCGGGCAAACAGTCGTTGCTGATTGGCGTTGCCACCTCCAGTCTGGCCCTGCACGCGCCGTCGCAAATTGTCGCGGCGATTAAATCTCGCGC
CGATCAACTGGGTGCCAGCGTGGTGGTGTCGATGGTAGAACGAAGCGGCGTCGAAGCCTGTAAAGCGGCGGTGCACAATCTTCTCGCGCAACGCGTCAGTGGGCTGATCATTAACTAT
CCGCTGGATGACCAGGATGCCATTGCTGTGGAAGCTGCCTGCACTAATGTTCCGGCGTTATTTCTTGATGTCTCTGACCAGACACCCATCAACAGTATTATTTACTCCCATGAGGACGGTA
CGCGACTGGGCGTGGAGCATCTGGTCGCATTGGGTCACCAGCAAATCGCGCTGTTAGCGGGCCCATTAAGTTCTGTCTCGGCGCGTCTGCGTCTGGCTGGCTGGCATAAATATCTCACT
CGCAATCAAATTCAGCCGATAGCGGAACGGGAAGGCGACTGGAGTGCCATGTCCGGTTTTCAACAAACCATGCAAATGCTGAATGAGGGCATCGTTCCCACTGCGATGCTGGTTGCCAA
CGATCAGATGGCGCTGGGCGCAATGCGCGCCATTACCGAGTCCGGGCTGCGCGTTGGTGCGGATATCTCGGTAGTGGGATACGACGATACCGAAGATAGCTCATGTTATATCCCGCCG
TTAACCACCATCAAACAGGATTTTCGCCTGCTGGGGCAAACCAGCGTGGACCGCTTGCTGCAACTCTCTCAGGGCCAGGCGGTGAAGGGCAATCAGCTGTTGCCAGTCTCACTGGTGA
AAAGAAAAACCACCCTGGCGCCCAATACGCAAACCGCCTCTCCCCGCGCGTTGGCCGATTCATTAATGCAGCTGGCACGACAGGTTTCCCGACTGGAAAGCGGGCAGTGATAAGGAT
CCTAATTGGTAACGAATCAGACAATTGACGGCTCGAGGGAGTAGCATAGGGTTTGCAGAATCCCTGCTTCGTCCATTTGACAGGCACATTATGCATCGATGATAAGCTGTCAAACATGAGC
AGATCCTCTACGCCGGACGCATCGTGGCCGGCATCACCGGCGCCACAGGTGCGGTTGCTGGCGCCTATATCGCCGACATCACCGATGGGGAAGATCGGGCTCGCCACTTCGGGCTC
ATGAGCAAATATTTTATCTGGCTTAACGATCGTTGGCTGTGTTGACAATTAATCATCGGCTCGTATAATGTGTGGAATTGTGAGCGCTCACAATTAGCTGTCACCGGATGTGCTTTCCGGTCTG
ATGAGTCCGTGAGGACGAAACAGCCTCTACAAATAATTTTGTTTAATACTAGAGAAAGAGGGGAAATACTAGATGATTATCGCGACTGCCGGACATGTGGATCATGGAAAGACAACATTGTT
GCAGGCGATTACTGGCGTAAATGCTGACCGTCTGCCGGAAGAAAAAAAGCGCGGCATGACCATAGATCTCGGCTATGCCTACTGGCCGCAGCCGGATGGTCGCGTGCCTGGTTTTATC
GACGTTCCCGGTCATGAAAAGTTTCTTTCCAACATGCTGGCGGGCGTTGGTGGTATCGATCACGCGCTGTTGGTGGTGGCATGCGATGACGGCGTGATGGCACAGACCCGTGAGCATCT
GGCGGTTTTGCAGCTGACCGGTAACCCGATGCTGACAGTGGCGCTGACCAAAGCCGATCGCGTGGACGAAGCGCGTGTTGATGAGGTTGAACGCCAGGTAAAGGAGGTTCTGCGGGA
ATACGGTTTTGCTGAGGCAAAACTGTTTATCACCGCAGCAACCGAAGGTCGGGGAATGGATGCCCTGCGCGAGCATCTGCTTCAGTTGCCGGATCGCGAGCACGCCAGCCAACATAGT
TTCCGCCTCGCGATTGACCGCGCATTTACCGTAAAAGGTGCCGGCCTGGTCGTCACCGGTACGGCGTTAAGCGGGGAAGTGAAGGTAGGCGATCCACTCTGGCTGACTGGTGTAAATA
AACCGATGCGTGTACGTGCGCTGCATGCGCAAAACCAGCCAACAGAAACCGCCAATGCCGGGCAGCGTATCGCGCTTAACATCGCGGGTGATGCGAAAAAAGAGCAGATTAACCGTG
GCGACTGGCTGCTTGCCGATGTGCCGCCAGAGCCGTTCACACGGGTGATTGTCGAGCTTCAAACCCATACATCGCTGACCCAGTGGCAGCCGCTGCATATTCACCACGCCGCCAGCC
ACGTCACGGGACGCGTTTCACTGCTGGAAGATAACCTTGCTGAACTGGTCTTCGACACCCCGTTATGGCTGGCAGATAACGACCGCCTGGTATTGCGCGATATCTCTGCCCGCAACACG
CTGGCCGGAGCGCGCGTCGTGATGCTTAACCCGCCGCGTCGCGGTAAACGTAAGCCGGAATATCTGCAATGGCTGGCGTCACTTGCACGGGCACAGAGCGATGCCGATGCGTTATCT
GTTCATCTGGAACGCGGCGCGGTTAACCTTGCGGATTTCGCCTGGGCGCGCCAGCTCAACGGCGAAGGGATGCGCGAATTGCTGCAACAGCCTGGTTATATTCAGGCTGGTTATAGCTT
GTTGAATACGCCGGTTGCCGCCCGCTGGCAGCGGAAAATTCTCGACACATTAGCGACTTATCATGAGCAACATCGCGATGAACCTGGCCCTGGGCGCGAACGTCTGCGACGTATGGCG
TTGCCAATGGAAGATGAAGCGCTGGTACTGTTGCTGATTAAAAAGATGCGCGAAAGCGGCGACATCCACAACCATCACGGCCGGCTGCATCTGCCAGATCACAAAGCGGGCCTCTAAC
ATTGTCTTATAAATACCAAAAATTTCACGTCTCTTGTTTAAAGAGGCGTGCTTTAATATTTTCTTAGAGTCCAATACATGAGAGTTTTAATTTCGTTTTTTTCCTTATTTTTTCTATTAGCGTTAGAGCT
GACTGTGGCCTTGATAGATCTTCGAAGCTTGGGCCCGAACAAAAACTCATCTCAGAAGAGGAGAGATAAATTGCACTGAAATCTAGAGTAACGGAATAGCTGTTCGTTGACTTGATAGACC
GATTGATTCATCATCTCATAAATAAAGAAAAACCACCGCTACCAACGGTGGTTTTCTCAAGGTTCGCTGAGCTACCAACTCTTTGAACCAAGGTAAGTGGGTTGGAGGACCGCACTCACCA
AAATCTGTTCTTTCAGTTTAGCCTTAACAGGTGCATAACTTCAAGACAAAGTCCTCTAAATCAGTTACCAATGGCTGCTGCCAGTGGCGATAAGTCGTGTCTTACCGGGTTGGACTCAAGAC
GATAGTTACCGGATAAGGCGCAGCGGTCGGGCTGAACGGGGGGTTCGTGCACACAGCCCAGCTTGGAGCGAACGACCTACACCGAACTGAGATACCAACAGCGTGAGCTATGAGAAA
GCGCCACGCTTCCCGAAGGGAGAAAGGCGGACAGGTATCCGGTAAGCGGCAGGGTCGGAACAGGAGAGCGCACGAGGGAGCTTCCAGGGGGAAACGCCTGGTATCTTTATAGTCCT
GTCGGGTTTCGCCACCTCTGGCTTGAGCGTCGATTTTTGTGATGCTCGTCAGGGGGGCGGAGCCTATGGAAAAACGCCTGCGGCGTTGGCTTCTTCCGGTGCTTTGCTTTTTGCTCACAT
GTTCTTTCCGGCTTTATCCCCTGATTCTGTGGATAACCGTATTACCGCTTTTGAGTGAGCTGACACCGCTCGCCGCAGTCGAACGACCGAGCGTAGCGAGTCAGTGAGCGAGGAAGCGG
AAGAGCGCTGCATGCCTATTTGTTTATTTTTCTAAATACATTCAAATATGTATCCGCTCATGAGACAATAACCCTGATAAATGCTTCAATAATATTGAAAAAGGAAGAGTATGAGCCATATTCAAC
GGGAAACGTCTTGCTCTAGGCCGCGATTAAATTCCAACATGGATGCTGATTTATATGGGTATAAATGGGCTCGCGATAATGTCGGGCAATCAGGTGCGACAATCTATCGATTGTATGGGAAG
CCCGATGCGCCAGAGTTGTTTCTGAAACATGGCAAAGGTAGCGTTGCCAATGATGTTACAGATGAGATGGTCAGACTAAACTGGCTGACGGAATTTATGCCTCTTCCGACCATCAAGCATT
TTATCCGTACTCCTGATGATGCATGGTTACTCACCACTGCGATCCCCGGGAAAACAGCATTCCAGGTATTAGAAGAATATCCTGATTCAGGTGAAAATATTGTTGATGCGCTGGCAGTGTTC
CTGCGCCGGTTGCATTCGATTCCTGTTTGTAATTGTCCTTTTAACAGCGACCGCGTATTTCGTCTCGCTCAGGCGCAATCACGAATGAATAACGGTTTGGTTGATGCGAGTGATTTTGATGAC
GAGCGTAATGGCTGGCCTGTTGAACAAGTCTGGAAAGAAATGCATAAACTTTTGCCATTCTCACCGGATTCAGTCGTCACTCATGGTGATTTCTCACTTGATAACCTTATTTTTGACGAGGGG
AAATTAATAGGTTGTATTGATGTTGGACGAGTCGGAATCGCAGACCGATACCAGGATCTTGCCATCCTATGGAACTGCCTCGGTGAGTTTTCTCCTTCATTACAGAAACGGCTTTTTCAAAAAT
ATGGTATTGATAATCCTGATATGAATAAATTGCAGTTTCATTTGATGCTCGATGAGTTTTTCTAACTGCAATAAGGTTGTTTTGCCGTGGTCAACGTGTCCGGCAGTCGCAATAATCATGGAGAA
GGGCGTTTAACATAACCACGGATTGTAACGTGAGATGGGTCAGGAGGACATATCGCGCATCAAGCCTTTGGCGGTTCGGTGCATCCCATGGCGATTGAATATTGAGCAGTAGAAAATATCT
GGATTGACAGTATTACAAGGGGTTGGAGAACGTCAGGAGAGGCATTTTGGCGGAAGATCACAGGAGTCGAACCTGCCCGGGACCGCTGGCGGCCCCAACTGGATTTAGAGTCCAGCC
GCCTCACCGGAGACGACGATCTTCCGCGCCTCGATTGCTACATGGAGGCGGGGCGCATTATAGCTACTTCCTTGAGTTTCTACATCCCCCAGATCAGTTTTTCCGCCTAATACACGCAG
CGTATTATCACGAAAGCATTATAGTTCATATAAAACCCATATTTTACAATGATATATGAGCGGAATTAACGCTACCACGATCACATAAAACAAGAATCATAAATTAATAACCAGATATCGGAATAT
TCGCTCTCACAGGGATGGCGGCCGC

**Supplementary Table 5| Plasmid sequence of plasmid recovered after 4 rounds of initial selection**

The ORF of NMC-A is pink, with the first TAG bolded; this was originally TGT. The ORF of SelB is green, the ORF of SelA is orange, and SelC is cyan, with the anticodon underlined. Duplicated genes are shown in white letters. See also **Supplementary Figure 2**.

AAAAAAAAACCACCGCTACCAACAGTGGTTTTCTCAAGGTTCGCTGAGCTACCAACTCTTTGAACCAAGGTAAGTGGGTTGGAGGACCGCACTCACCAAAATCTGTTCTTTCAGTTTAGCCTTAACAGGTG
CATAACTTCAAGACAAAGTCCTCTAAATCAGTTACCAATGGCTGCTGCCAGTGGCGATAAGTCGTGTCTTACCGGGTTGGACTCAAGACGATAGTTACCGGATAAGGCGCAGCGGTCGGGCTGAACGGG
GGGTTCGTGCACACAGCCCAGCTTGGAGCGAACGACCTACACCGAACTGAGATACCAACAGCGTGAGCTATGAGAAAGCGCCACGCTTCCCGAAGGGAGAAAGGCGGACAGGTATCCGGTAAGCGGC
AGGGTCGGAACAGGAGAGCGCACGAGGGAGCTTCCAGGGGGAAACGCCTGGTATCTTTATAGTCCTGTCGGGTTTCGCCACCTCTGGCTTGAGCGTCGATTTTTGTGATGCTCGTCAGGGGGGCGGAG
CCTATGGAAAAACGCCTGCGGCGTTGGCTTCTTCCGGTGCTTTGCTTTTTGCTCACATGTTCTTTCCGGCTTTATCCCCTGATTCTGTGGATAACCGTATTACCGCTTTTGAGTGAGCTGACACCGCTCGCCG
CAGTCGAACGACCGAGCGTAGCGAGTCAGTGAGCGAGGAAGCGGAAGAGCGCTGCATGCCTATTTGTTTATTTTTCTAAATACATTCAAATATGTATCCGCTCATGAGACAATAACCCTGATAAATGCTT
CAATAATATTGAAAAAGGAAGAGTATGAGCCATATTCAACGGGAAACGTCTTGCTCTAGGCCGCGATTAAATTCCAACATGGATGCTGATTTATATGGGTATAAATGGGCTCGCGATAATGTCGGGCAAT
CAGGTGCGACAATCTATCGATTGTATGGGAAGCCCGATGCGCCAGAGTTGTTTCTGAAACATGGCAAAGGTAGCGTTGCCAATGATGTTACAGATGAGATGGTCAGACTAAACTGGCTGACGGAATTTA
TGCCTCTTCCGACCATCAAGCATTTTATCCGTACTCCTGATGATGCATGGTTACTCACCACTGCGATCCCCGGGAAAACAGCATTCCAGGTATTAGAAGAATATCCTGATTCAGGTGAAAATATTGTTGATG
CGCTGGCAGTGTTCCTGCGCCGGTTGCATTCGATTCCTGTTTGTAATTGTCCTTTTAACAGCGACCGCGTATTTCGTCTCGCTCAGGCGCAATCACGAATGAATAACGGTTTGGTTGATGCGAGTGATTTTG
ATGACGAGCGTAATGGCTGGCCTGTTGAACAAGTCTGGAAAGAAATGCATAAACTTTTGCCATTCTCACCGGATTCAGTCGTCACTCATGGTGATTTCTCACTTGATAACCTTATTTTTGACGAGGGGAAA
TTAATAGGTTGTATTGATGTTGGACGAGTCGGAATCGCAGACCGATACCAGGATCTTGCCATCCTATGGAACTGCCTCGGTGAGTTTTCTCCTTCATTACAGAAACGGCTTTTTCAAAAATATGGTATTGAT
AATCCTGATATGAATAAATTGCAGTTTCATTTGATGCTCGATGAGTTTTTCTAACTGCAATAAGGTTGTTTTGCCGTGGTCAACGTGTCCGGCAGTCGCAATAATCATTTCAACAACATCTCCAAAAACCGTT
GCTCATCTTCAAGGCAGCGTAAATCCAGCCACAATCGTCCGTCATAAATACGACCAATCACCGGCACTGGCAATTCACGCCAGCGAGCGGCTAATGACTCAAGGTGGCTACCGCGTCCATCATGGGGTGT
AAACGTTAATGCCGCGCTCGGCAGGCGATCAACCGGCAGCGAACCACTGCCAATCTGCGAAAGACATGGCATAACCTGTACCGCAAACTCCGCGCCGTAATGTGCGGCAAGGGGGGCCTGTAAACGTT
GTGCCTGGATTTGAATGACCTCTGCGCTGCGGGTAAGCAGGCGCAGGGTCGGTAATTTTTCACTCAGAGCTTCAGGGTGTAAATAAAGACGCAACGTGGCTTCCAGCGCCGCGAGGGTCATTTTATCCGC
GCGTAATGCACGCTTCAGCGGGTGGCTTTGCAGGCGGGCGATCATCTCTTTTTTACCAACAATAATTCCTGCCTGCGGCCCGCCTAACAACTTGTCGCCGGAGAAACTCACCAGACTGACGCCCGCCGCAA
TCAACTCCTGCGGCATTGGCTCTTTCGGCAAACCGTACTGGCTAAGATCGACCAGCGAGCCACTGCCTAAATCAGTCACTACGGGAACATCCAGCTCTTTGCCGAGCGCCACCAGTTCCGCTTCATCTATC
GCTTTGGTGAACCCCTGAATGCTGTAGTTACTGGTATGTACTTTCATCAACAGTGCGGTATTTTCATTCACCGCCTGACGATAATCATTCGCGTGCGTGCGGTTGGTGGTCCCTACTTCGTGTAGGGTGCAG
CCTGCCTGACGCATAACATCGGGAATACGAAACGCGCCGCCAATCTCCACCAGTTCGCCGCGAGATACCACCACCTCTTTTCCGCTGGCAGTGGCCGCCAACATCAATAACACCGCCGCCGCATTGTTATT
GACGATACAGGCATCTTCCGCCCCCGTAATACGGCACAGCAGCTGCGCCAGCGCCCGATCGCGATGTCCGCGTCCGGCGTCGTCCAGATCATACTCGAGGGTCACTGGCGAACGCATAGCCTGCGCAAC
GGCTTCCACCGCGGCTTCCGCCTGTAAAGCTCGCCCAAGGTTGGTATGCAGCACGGTTCCCGTCAGGTTGATCACCGGACGCAGCGCGCTCTGCGCTTCTTTCGTCAACCGGGCATCGACTTCTTGCGCCC
AGTTTTCACACCACGCAGGCAGCGTCTGGCTGCCACGAATCACTTCTCGCGCTTCGTCGAGCATCTGACGCAACAATTCCACCACGCGGGTGTGACCATAAGTATCACGCAAAGAAAGGAAGGAGCTATC
GCGCAATAAGCGATCAATAGCCGGAAGTTGACTATAGAGGAAACGCGTTTCGGTTGTCATGGTTTAGTTCCTCACCTTGTCGTATTATACTATGCCGATATACTATGCCGATGATTAGGAAACCTGGCTGA
TCAAGGCCCTCTCACACGGAGAAGGGCGTTTAACATAACCACGGATTGTAACGTGAGATGGGTCAGGAGGACATATCGCGCATCAAGCCTTTGGCGGTTCGGTGCATCCCATGGCGATTGAATATTGAG
CAGTAGAAAATATCTGGATTGACAGTATTACAAGGGGTTGGAGAACGTCAGGAGAGGCATTTTGGCGGAAGATCACAGGAGTCGAACCTGCCCGGGACCGCTGGCGGCCCCAACTGGATTTAGAGTCC
AGCCGCCTCACCGGAGACGACGATCTTCCGCGCCTCGATTGCTACATGGAGGCGGGGCGCATTATAGCTACTTCCTTGAGTTTCTACATCCCCCAGATCAGTTTTTTCCGCCTAATACACGCAGCGTATTAT
CACGAAAGCATTATAGTTCATATAAAACCCATATTTTACAATGATATATGAGCGGAATTAACGCTACCACGATCACATAAAACAAGAATCATAAATTAATAACCAGATATCGGAATATTCGCTCTCACAGG
GATGGCGGCCGCTCGACCGATGCCCTTGAGACGTTGATCGGCACGTATGGCAGATAGTAAATTTTATAGATTTAAATCAACATCTATAACTGTATAATTCGTTTTCTCAACTCATTACAACACTCCGTTAGT
AATGAAGCTCATTCTATACAATGACAATTAATAGGTAAAGTTATGTCACTTAATGTAAAGCAAAGTAGAATAGCCATCTTGTTTAGCTCTTGTTTAATTTCAATATCATTTTTCTCACAGGCCAATACGAAGG
GCATTGATGAGATTAAAAACCTTGAAACAGATTTCAATGGCAGGATTGGTGTCTACGCTTTAGACACTGGCTCGGGTAAATCATTTTCGTACAGAGCAAATGAACGATTTCCATTATAGAGTTCTTTTAAA
GGTTTTTTAGCTGCTGCTGTATTAAAAGGCTCTCAAGATAATCGACTTAATCTTAATCAGATTGTGAATTATAATACAAGAAGTTTAGAGTTCCATTCACCCATCACAACTAAATATAAAGATAATGGAATG
TCATTAGGTGATATGGCTGCTGCTGCTTTACAATATAGCGACAATGGTGCTACTAATATTATTCTTGAACGTTATATCGGTGGTCCAGAGGGTATGACTAAATTCATGCGGTCGATTGGAGATGAAGATTT
TAGACTCGATCGTTGGGAGTTAGATCTAAACACAGCTATTCCAGGCGATGAGCGTGACACATCTACACCTGCAGCAGTAGCCAAGAGTCTGAAAACCCTTGCTCTGGGTAACATACTTAGTGAACATGAA
AAGGAAACCTATCAGACATGGTTAAAGGGTAACACAACCGGTGCAGCGCGTATTCGTGCTAGCGTACCAAGCGATTGGGTAGTTGGCGATAAAACTGGTAGTTGCGGAGCATACGGTACGGCAAATGA
TTATGCGGTAGTCTGGCCAAAGAACCGGGCTCCTCTTATAATTTCTGTATACACAACAAAAAACGAAAAAGAAGCCAAGCATGAGGATAAAGTAATCGCAGAAGCTTCAAGAATTGCAATTGATAACCTT
AAATAACAAAGCCCGCCGAAAGGCGGGCTTTTTTTTGGATCCTGTAGAAACGCAAAAAGGCCATCCGCATGACTAACATGAGAATTACAACTATCACCGCAAGGGATAAATATCTAACACCGTTATGAC
AACTTGACGGCTACATCATTCACTTTTTCTTCACAACCGGCACGGAACTCGCTCGGGCTGGCCCCGGTGCATTTTTTAAATACCCGCGAGAAGTAGAGTTGATCGTCAAAACCAACATTGCGACCGACGGT
GGCGATAGGCATCCGGGTGGTGCTCAAAAGCAGCTTCGCCTGGCTGATACGTTGGTCCTCGCGCCAGCTTAAGACGCTAATCCCTAACTGCTGGCGGAAAAGATGTGACAGACGCGACGGCGACAAGC
AAACATGCTGTGCGACGCTGGCGATATCAAAATTGCTGTCTGCCAGGTGATCGCTGATGTACTGACAAGCCTCGCGTACCCGATTATCCATCGGTGGATGGAGCGACTCGTTAATCGCTTCCATGTGCCG
CAGTAACAATTGCTCAAGCAGATTTATCGCCAGCAGCTCCGAATAGCGCCCTTCCCCTTGCCCGGCGTTAATGATTTGCCCAAACAGGTCGCTGAAATGCGGCTGGTGCGCTTCATCCGGGCGAAAGAAC
CCCGTATTGGCAAATATTGACGGCCAGTTAAGCCATTCATGCCAGTAGGCGCGCGGACGAAAGTAAACCCACTGGTGATACCATTCGCGAGCCTCCGGATGACGACCGTAGTGATGAATCTCTCCTGGCG
GGTTTAACGCCGATTGAGGCCAACGGTTATCTCGATTTTTTTATCGACCGACCGCTGGGAATGAAAGGTTATATTCTCAATCTCACCATTCGCGGTCAGGGGGTGGTGAAAAATCAGGGACGAGAATTTG
TTTGCCGACCGGGTGATATTTTGCTGTTCCCGCCAGGAGAGATTCATCACTACGGTCGTCATCCGGAGGCTCGCGAATGGTATCACCAGTGGGTTTACTTTCGTCCGCGCGCCTACTGGCATGAATGGCTT
AACTGGCCGTCAATATTTGCCAATACGGGGTTCTTTCGCCCGGATGAAGCGCACCAGCCGCATTTCAGCGACCTGTTTGGGCAAATCATTAACGCCGGGCAAGGGGAAGGGCGCTATTCGGAGCTGCTG
GCGATAAATCTGCTTGAGCAATTGTTACTGCGGCACATGGAAGCGATTAACGAGTCGCTCCATCCACCGATGGATAATCGGGTACGCGAGGCTTGTCAGTACATCAGCGATCACCTGGCAGACAGCAATT
TTGATATCGCCAGCGTCGCACAGCATGTTTGCTTGTCGCCGTCGCGTCTGTCACATCTTTTCCGCCAGCAGTTAGGGATTAGCGTCTTAAGCTGGCGCGAGGACCAACGTATCAGCCAGGCGAAGCTGCTT
TTGAGCACCACCCGGATGCCTATCGCCACCGTCGGTCGCAATGTTGGTTTTGACGATCAACTCTACTTCTCGCGGGTATTTAAAAAATGCACCGGGGCCAGCCCGAGCGAGTTCCGTGCCGGTTGTGAAG
AAAAAGTGAATGATGTAGCCGTCAAGTTGTCATAACGGTGTTAGATATTTATCCCTTGCGGTGATAGTTGTAATTCTCATGTTAGTCATGCGGATGGCCTTTTTGCGTTTCTACAGGATCCAAAAAAAGCCC
GCCTTTCGGCGGGCTTTGTTATTTAAGGTTATCAATTGCAATTCTTGAAGCTTCTGCGATTACTTTATCCTCATGCTTGGCTTCTTTTTCGTTTTTTGTTGTGTATACAGAAATTATAAGAGGAGCCCGGTTCT
TTGGCCAGACTACCGCATAATCATTTGCCGTACCGTATGCTCCGCAACTACCAGTTTTATCGCCAACTACCCAATCGCTTGGTACGCTAGCACGAATACGCGCTGCACCGGTTGTGTTACCCTTTAACCATG
TCTGATAGGTTTCCTTTTCATGTTCACTAAGTATGTTACCCAGAGCAAGGGTTTTCAGACTCTTGGCTACTGCTGCAGGTGTAGATGTGTCACGCTCATCGCCTGGAATAGCTGTGTTTAGATCTAACTCCC
AACGATCGAGTCTAAAATCTTCATCTCCAATCGACCGCATGAATTTAGTCATACCCTCTGGACCACCGATATAACGTTCAAGAATAATATTAGTAGCACCATTGTCGCTATATTGTAAAGCAGCAGCAGCCA
TATCACCTAATGACATTCCATTATCTTTATATTTAGTTGTGATGGGTGAATGGAACTCTAAACTTCTTGTATTATAATTCACAATCTGATTAAGATTAAGTCGATTATCTTGAGAGCCTTTTAATACAGCAGC
AGCTAAAAAACCTTTAAAAGAACTCTATAATGGAAATCGTTCATTTGCTCTGTACGAAAATGATTTACCCGAGCCAGTGTCTAAAGCGTAGACACCAATCCTGCCATTGAAATCTGTTTCAAGGTTTTTAAT
CTCATCAATGCCCTTCGTATTGGCCTGTGAGAAAAATGATATTGAAATTAAACAAGAGCTAAACAAGATGGCTATTCTACTTTGCTTTACATTAAGTGACATAACTTTACCTATTAATTGTCATTGTATAGAA
TGAGCTTCATTACTAACGGAGTGTTGTAATGAGTTGAGAAAACGAATTATACAGTTATAGATGTTGATTTAAATCTATAAAATTTACTATCTGCCATACGTGCCGATCAACGTCTCAAGGGCATCGGTCGA
GCGGCCGCCATCCCTGTGAGAGCGAATATTCCGATATCTGGTTATTAATTTATGATTCTTGTTTTATGTGATCGTGGTAGCGTTAATTCCGCTCATATATCATTGTAAAATATGGGTTTTATATGAACTATAA
TGCTTTCGTGATAATACGCTGCGTGTATTAGGCGGAAAAAACTGATCTGGGGGATGTAGAAACTCAAGGAAGTAGCTATAATGCGCCCCGCCTCCATGTAGCAATCGAGGCGCGGAAGATCGTCGTCTC
CGGTGAGGCGGCTGGACTCTAAATCCAGTTGGGGCCGCCAGCGGTCCCGGGCAGGTTCGACTCCTGTGATCTTCCGCCAAAATGCCTCTCCTGACGTTCTCCAACCCCTTGTAATACTGTCAATCCAGATA
TTTTCTACTGCTCAATATTCAATCGCCATGGGATGCACCGAACCGCCAAAGGCTTGATGCGCGATATGTCCTCCTGACCCATCTCACGTTACAATCCGTGGTTATGTTAAACGCCCTTCTCCGTGTGAGAGG
GCCTTGATCAGCCAGGTTTCCTAATCATCGGCATAGTATATCGGCATAGTATAATACGACAAGGTGAGGAACTAAACCATGACAACCGAAACGCGTTTCCTCTATAGTCAACTTCCGGCTATTGATCGCTT
ATTGCGCGATAGCTCCTTCCTTTCTTTGCGTGATACTTATGGTCACACCCGCGTGGTGGAATTGTTGCGTCAGATGCTCGACGAAGCGCGAGAAGTGATTCGTGGCAGCCAGACGCTGCCTGCGTGGTGT
GAAAACTGGGCGCAAGAAGTCGATGCCCGGTTGACGAAAGAAGCGCAGAGCGCGCTGCGTCCGGTGATCAACCTGACGGGAACCGTGCTGCATACCAACCTTGGGCGAGCTTTACAGGCGGAAGCCG
CGGTGGAAGCCGTTGCGCAGGCTATGCGTTCGCCAGTGACCCTCGAGTATGATCTGGACGACGCCGGACGCGGACATCGCGATCGGGCGCTGGCGCAGCTGCTGTGCCGTATTACGGGGGCGGAAGAT
GCCTGTATCGTCAATAACAATGCGGCGGCGGTGTTATTGATGTTGGCGGCCACTGCCAGCGGAAAAGAGGTGGTGGTATCTCGCGGCGAACTGGTGGAGATTGGCGGCGCGTTTCGTATTCCCGATGTT
ATGCGTCAGGCAGGCTGCACCCTACACGAAGTAGGGACCACCAACCGCACGCACGCGAATGATTATCGTCAGGCGGTGAATGAAAATACCGCACTGTTGATGAAAGTACATACCAGTAACTACAGCATT
CAGGGGTTCACCAAAGCGATAGATGAAGCGGAACTGGTGGCGCTCGGCAAAGAGCTGGATGTTCCCGTAGTGACTGATTTAGGCAGTGGCTCGCTGGTCGATCTTAGCCAGTACGGTTTGCCGAAAGA
GCCAATGCCGCAGGAGTTGATTGCGGCGGGCGTCAGTCTGGTGAGTTTCTCCGGCGACAAGTTGTTAGGCGGCCCGCAGGCAGGAATTATTGTTGGTAAAAAAGAGATGATCGCCCGCCTGCAAAGCC
ACCCGCTGAAGCGTGCATTACGCGCGGATAAAATGACCCTCGCGGCGCTGGAAGCCACGTTGCGTCTTTATTTACACCCTGAAGCTCTGAGTGAAAAATTACCGACCCTGCGCCTGCTTACCCGCAGCGC
AGAGGTCATTCAAATCCAGGCACAACGTTTACAGGCCCCCCTTGCCGCACATTACGGCGCGGAGTTTGCGGTACAGGTTATGCCATGTCTTTCGCAGATTGGCAGTGGTTCGCTGCCGGTTGATCGCCTG
CCGAGCGCGGCATTAACGTTTACACCCCATGATGGACGCGGTAGCCACCTTGAGTCATTAGCCGCTCGCTGGCGTGAATTGCCAGTGCCGGTGATTGGTCGTATTTATGACGGACGATTGTGGCTGGATT
TACGCTGCCTTGAAGATGAGCAACGGTTTTTGGAGGATTGTATGATTATCGCGACTGCCGGACATGTGGATCATGGAAAGACAACATTGTTGCAGGCGATTACTGGCGTAAATGCTGACCGTCTGCCGGA
AGAAAAAAAGCGCGGCATGACCATAGATCTCGGCTATGCCTACTGGCCGCAGCCGGATGGTCGCGTGCCTGGTTTTATCGACGTTCCCGGTCATGAAAAGTTTCTTTCCAACATGCTGGCGGGCGTTGGT
GGTATCGATCACGCGCTGTTGGTGGTGGCATGCGATGACGGCGTGATGGCACAGACCCGTGAGCATCTGGCGGTTTTGCAGCTGACCGGTAACCCGATGCTGACAGTGGCGCTGACCAAAGCCGATCG
CGTGGACGAAGCGCGTGTTGATGAGGTTGAACGCCAGGTAAAGGAGGTTCTGCGGGAATACGGTTTTGCTGAGGCAAAACTGTTTATCACCGCAGCAACCGAAGGTCGGGGAATGGATGCCCTGCGCG
AGCATCTGCTTCAGTTGCCGGATCGCGAGCACGCCAGCCAACATAGTTTCCGCCTCGCGATTGACCGCGCATTTACCGTAAAAGGTGCCGGCCTGGTCGTCACCGGTACGGCGTTAAGCGGGGAAGTGA
AGGTAGGCGATCCACTCTGGCTGACTGGTGTAAATAAACCGATGCGTGTACGTGCGCTGCATGCGCAAAACCAGCCAACAGAAACCGCCAATGCCGGGCAGCGTATCGCGCTTAACATCGCGGGTGATG
CGGAAAAAGAGCAGATTAACCGTGGCGACTGGCTGCTTGCCGATGTGCCGCCAGAGCCGTTCACACGGGTGATTGTCGAGCTTCAAACCCATACATCGCTGACCCAGTGGCAGCCGCTGCATATTCACCA
CGCCGCCAGCCACGTCACGGGACGCGTTTCACTGCTGGAAGATAACCTTGCTGAACTGGTCTTCGACACCCCGTTATGGCTGGCAGATAACGACCGCCTGGTATTGCGCGATATCTCTGCCCGCAACACG
CTGGCCGGAGCGCGCGTCGTGATGCTTAACCCGCCGCGTCGCGGTAAACGTAAGCCGGAATATCTGCAATGGCTGGCGTCACTTGCACGGGCACAGAGCGATGCCGATGCGTTATCTGTTCATCTGGAA
CGCGGCGCGGTTAACCTTGCGGATTTCGCCTGGGCGCGCCAGCTCAACGGCGAAGGGATGCGCGAATTGCTGCAACAGCCTGGTTATATTCAGGCTGGTTATAGCTTGTTGAATGCGCCGGTTGCCGCC
CGCTGGCAGCGGAAAATTCTCGACACATTAGCGACTTATCATGAGCAACATCGCGATGAACCTGGCCCTGGGCGCGAACGTCTGCGACGTATGGCGTTGCCAATGGAAGATGAAGCGCTGGTACTGTTG
CTGATTGAAAAGATGCGCGAAAGCGGCGACATCCACAACCATCACGGCCGGCTGCATCTGCCAGATCACAAAGCGGGCCTCTAACATTGTCTTATAAATACCAAAAATTTCACATCTCTTGTTTAAAGAG
GCGTGCTTTAATATTTTCTTAGAGTCCAATACATGAGAGTTTTAATTTCGTTTTTTTCCTTATTTTTTCTATTAGCGTTAGAGCTGACTGTGGCCTTGATAGATCTTCGAAGCTTGGGCCCGAACAAAAACTC
ATCTCAGAAGAGGATCTGAGAGATAAATTGCACTGAAATCTAGAGTAACGGAATAGCTGTTCGTTGACTTGATAGACCGATTGATTCATCATCTCATAAAT

**Supplementary Table 6| Oligonucleotide sequences used in this study**

| Name | Sequence (5'-3') |
|---|---|
| Oligo1 | CATACAATCCTCCTAACGGGT |
| Oligo2 | TAACATTGTCTTATAAATACCAAAAATTTCAC |
| Oligo3 | ACCCGTTAGGAGGATTGTATG |
| Oligo4 | GTGAAATTTTTGGTATTTATAAGACAATGTTA |
| Oligo5 | GCATGGTCTCTTACCCGTTAGGAGGATTGTATG |
| Oligo6 | GCATGGTCTCTGTGAAATTTTTGGTATTTATAAGACAATGTTA |
| Oligo7 | GCATGGTCTCTGGTATGGAGAAACAGTAGAGAG |
| Oligo8 | GCATGGTCTCTTCACGTCTCTTGTTTAAAGAGGC |
| Oligo9 | GCATGGTCTCTCTAGAGAAAGAGGGGAAATACTAGATGATCATCGCCACGGCGGGTCATGTGGATCATGGAAAGACAACA |
| Oligo10 | GCATGGTCTCTCTAGTATTAAACAAAATTATTTGTAGAGG |
| Oligo11 | GCCGAGATCTATGGTCATGC |
| Oligo12 | GCATGACCATAGATCTCGGCNNSGCCNNSTGGCCGCAGCCGGATGG |
| Oligo13 | GGCCGGCACCTTTTACGGTAAATGCSNNGTCAATCGCGAGGCGGAAA |
| Oligo14 | CCGTAAAAGGTGCCGGCCTGGTCGTCNNSGGTACGGCGTTAAGCGGG |
| Oligo15 | CTGCCCGGCATTGGCGGT |
| Oligo16 | AAACCGCCAATGCCGGGCAGNNSATCGCGCTTAACATCGCGG |
| Olugo17 | GCATGGTCTCTCTAGAGAAAGAGGGGAAATACTAG |

N denotes for all possible nucleotides and S denotes either C or G.

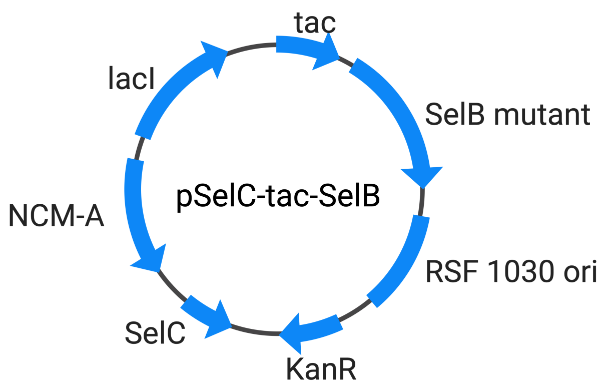


**Supplementary Figure 1| Plasmid map of pSelC-tac-SelB**

Compared to pSelAC-araBAD-SelB, SelA gene was removed as well as the promoter of SelB was changed from araBAD to tac-lac promoter. araC was replaced with LacI as well.

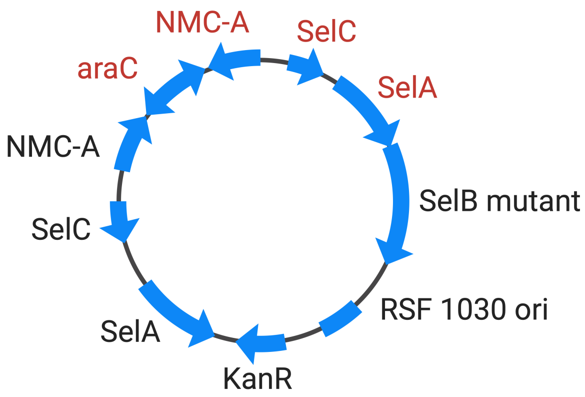


**Supplementary Figure 2| Plasmid map of SelB-v1-containing plasmid recovered after round 4.** After round 4, the plasmid backbone had duplications in NMC-A and SelC (highlighted with red letters), while SelB was in-frame with the duplicated SelA. The sequence is shown in **Supplementary Table 5**.

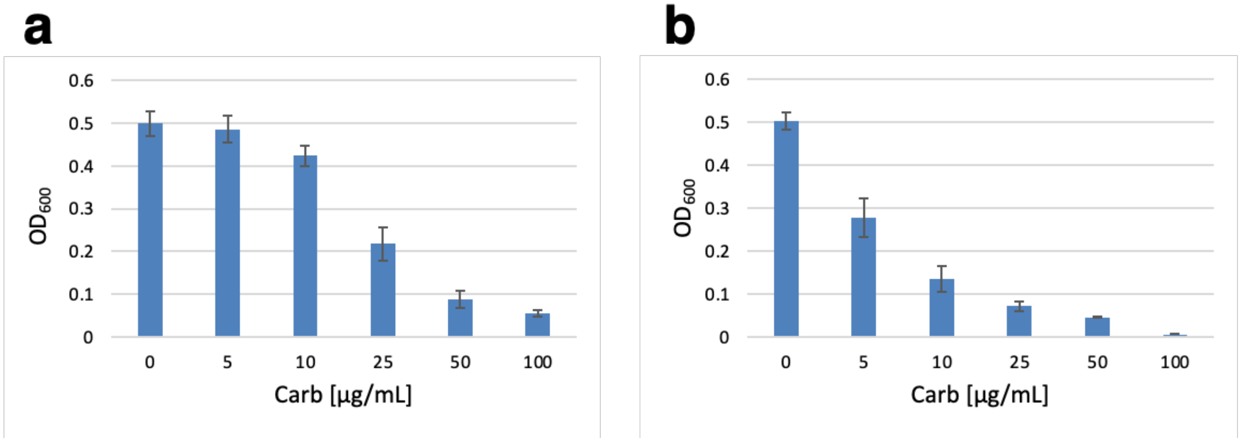


**Supplementary Figure 3| NMC-A assay of SelB-v1**

(a) NMC-A assay of SelB-v1 having one TAG codon in the NMC-A gene. (b) NMC-A assay of SelB-v1 using two TAG codons in the NMC-A gene.

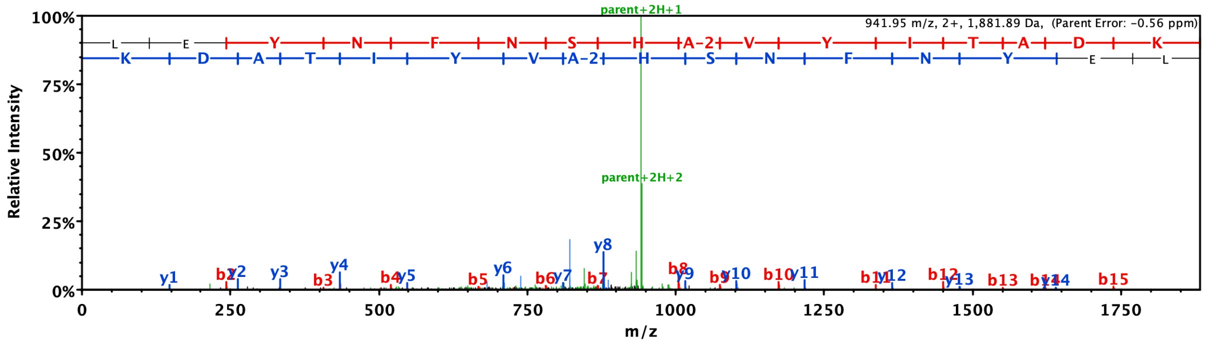


**Supplementary Figure 4| LC-MS result of trypsin-digested sfGFP expressed in the presence of selenium and SelB-v2**

Ion fragments corresponding to dehydroalanine incorporation were observed in LC-MS/MS. Shown is an example of the spectrum of a trypsin-digested peptide from sfGFP whose sequence is LEYNFNSHXVYITADK (m/z 1881.89), where X corresponds to dehydroalanine. Both b and y ions from this peptide fragments were observed, indicating dehydroalanine incorporation.

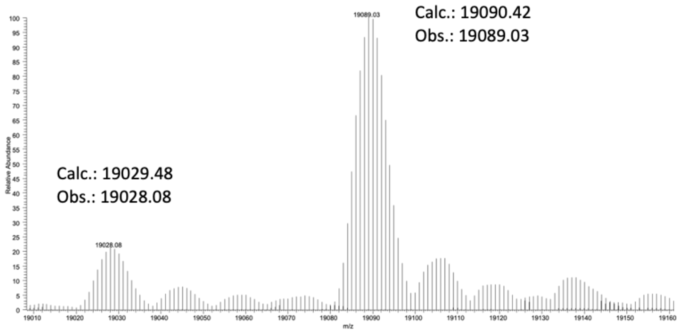


**Supplementary Figure 5| Mass spectrometry of DHFR P61TAG in a C321dSelABC strain background.**

The deconvoluted mass spectrum of purified DHRF S61TAG protein which was expressed in the presence of SelB-v2 in C321 dSelABC. Mass spectrometry shows that both serine-containing DHFR and selenocysteine-containing DHFR were present in the sample mixture: the calculated mass for DHRF P61S (serine-containing) is 19029.48 and the observed mass is 19028.08, while the calculated mass for DHFR P61U (selenocysteine-containing) is 19090.42 and the observed mass is 19089.03. Calc., calculated; Obs., observed.

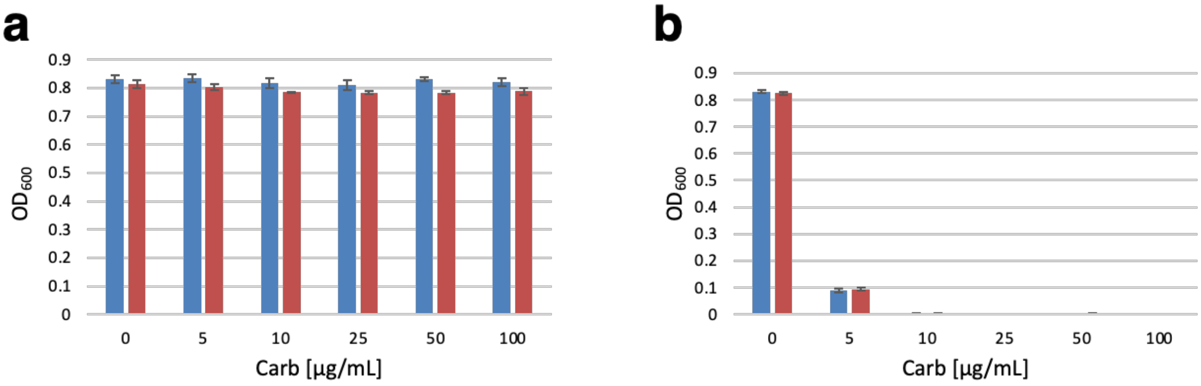


**Supplementary Figure 6| NMC-A assay with S71 mutants**

(a) NMC-A assay using wild-type NMC-A. The blue bar refers to growth without induction of SelB-v2, while the red bar indicates growth with induction with SelB-v2. Each data point represents the mean of three independent experiments ± s.d (standard deviation). (b) NMC-A assay of the NMC-A S71C mutant. The blue bar refers to growth without induction of SelB-v2, while the red bar indicates growth with induction with SelB-v2. Each data point represents the mean of three independent experiments ± s.d (standard deviation).

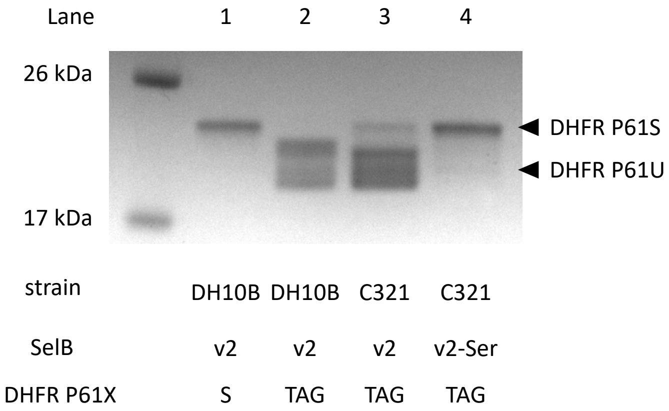


**Supplementary Figure 7| SDS-PAGE of purified DHFR expressed in different strains**

SDS-PAGE analysis of purified DHFR expressed in either the DH10B dSelABC strain or the C321 dSelABC strain. Lane 1 corresponds to purified DHFR P61S while lanes 2-4 are purified DHFR P61TAG variants. For lanes 1-3 SelB-v2 was induced, while for lane 4, SelB-v2-Ser was induced, instead of SelB-v2. For the DHFR P61TAG protein that was expressed in DH10B dSelABC with SelB-v2, only DHFR P61U (selenocysteine) was observed, while no DHFR P61S (serine) (lane 2). In the context of C321 dSelABC, however, some DHFR P61S (serine) was observed (lane 3). When DHFR P61TAG was expressed in the presence of SelB-v2-Ser, however, primarily DHFR P61S (serine) was observed (lane 4).

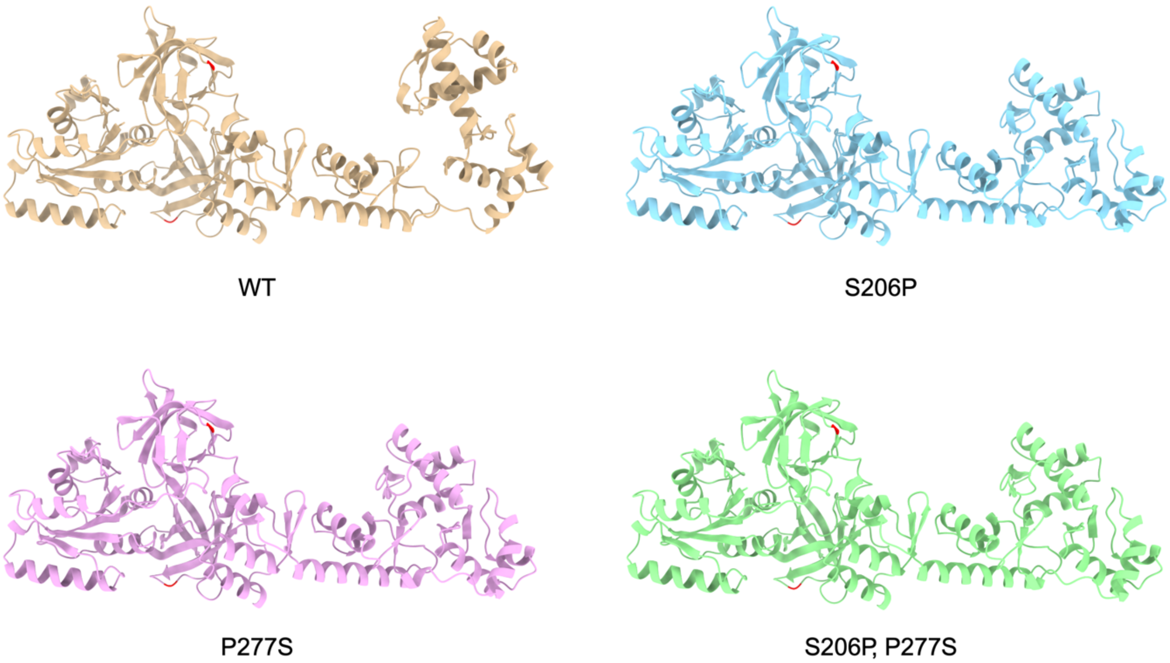


**Supplementary Figure 8| Non-truncated SelB RNA-binding domain undergoes conformational change upon S206P and P277S mutations.**

Structures generated with AlphaFold3, pTM scores of 0.81, 0.82, 0.82, 0.83 for WT, S206P, P277S, and [S206P, P277S] double mutant, respectively. Residues 206 and 277 are highlighted in red. The RNA-binding domain (right side) undergoes a large conformational change upon the introduction of these mutations.
